# DONOR VARIABILITY IN HUMAN MESENCHYMAL STEM CELL OSTEOGENIC RESPONSE AS A FUNCTION OF PASSAGE CONDITIONS AND DONOR SEX

**DOI:** 10.1101/2023.11.12.566781

**Authors:** Vasiliki Kolliopoulos, Aleczandria Tiffany, Maxwell Polanek, Brendan A.C. Harley

## Abstract

Contemporary tissue engineering efforts often seek to use mesenchymal stem cells (MSCs) due to their potential to differentiate to various tissue-specific cells and generate a pro-regenerative secretome. While MSC differentiation and therapeutic potential can differ as a function of matrix environment, it may also be widely influenced as a function of donor-to-donor variability. Further, effects of passage number and donor sex may further convolute the identification of clinically effective MSC-mediated regeneration technologies. We report efforts to adapt a well-defined mineralized collagen scaffold platform to study the influence of MSC proliferation and osteogenic potential as a function of passage number and donor sex. Mineralized collagen scaffolds broadly support MSC osteogenic differentiation and regenerative potency in the absence of traditional osteogenic supplements for a wide range of MSCs (rabbit, rat, porcine, human). We obtained a library of bone marrow and adipose tissue derived stem cells to examine donor-variability of regenerative potency in mineralized collagen scaffolds. MSCs displayed reduced proliferative capacity as a function of passage duration. Further, MSCs showed significant sex-based differences. Notably, MSCs from male donors displayed significantly higher metabolic activity and proliferation while MSCs from female donor displayed significantly higher osteogenic response via increased alkaline phosphate activity, osteoprotegerin release, and mineral formation in vitro. Our study highlights the essentiality of considering MSC donor sex and culture expansion in future studies of biomaterial regenerative potential.

## 1. Introduction

There were 178 million bone fractures that occurred globally in 2019 [1]. The clinical gold-standard for bone graft substitutes are allografts and autografts [2–5], with ∼2 million autologous or allogenic/artificial bone grafts used between 1992 and 2007 in the United States [6]. Both suffer from key limitations. Notably, limited donor tissue and the need to create a secondary injury site for autografts [2–5]. And concerns about inconsistent purification methods, infection or rejection [2–5, 7, 8], and variability in healing based on both donor and recipient biology exist for allografts [2–5]. Tissue engineering offers a framework to develop biomaterials for bone regeneration that may address limitations in allograft and autograft approaches. Many contemporary tissue engineering efforts seek to combine exogenous mesenchymal stem cells (MSCs), morphogens, and biomaterials for bone tissue repair [9–12]. MSCs are capable of both differentiating to tissue-specific lineages as well as generating a complex secretome to support the regenerative action of other cells in the wound site. Further, they can be derived from multiple tissues like adipose (fat tissue) and bone marrow [13].

Tissue engineering approaches often rely on cells of unknown origin or that lack diversity related to sex, age, and ethnicity [14, 15]. Indeed, bone healing can be influenced by many factors including pre-existing health conditions, age, and sex. Further, sex hormones are important during development of the skeletal system; hormonal differences can impact peak bone mass and microarchitecture of bone tissue [16–18] and may also underlie sex-differences in bone healing following injury [19, 20]. Broadly, male patients have more robust bone healing than female patients [19, 20]. Sex differences in bone healing are also observed by groups utilizing animal models, with male animals exhibiting more rapid and robust bone healing than female animals [21, 22]. Yet, a recent comprehensive review surveyed over 300 articles in the biomaterials literature from 2019 that used in vitro cell experiments, and found more than 95% of studies using cell lines, and approximately 90% of studies using primary cells, failed to report the sex of cells used [23]. Further, potential variability in healing capacity between donors (e.g., age, underlying health conditions) pose additional challenges for developing biomaterial implants to heal bone injuries [24–27]. Hence it is essential regenerative medicine studies more definitively identify the sex, ancestry, age, and underlying health conditions of the cells used.

Regenerative medicine technologies often require large numbers of MSCs, motivating studies to understand the confounding effect of *in vitro* expansion conditions. *In vitro* monolayer culture typically used for MSC expansion can significantly influence the phenotype, proliferation, and differentiation capacity of hMSCs [28–32]. Further, most studies on the influence of these factors utilize two-dimensional (2D) tissue culture plastic for their culture and subsequent analyses which is not representative of the 3D environment these cells would normally experience. Nonetheless, several groups have found that hMSC activity including differentiation capacity, proliferation, and expression profiles is influenced by donor age [31–34] and passage number [32, 35, 36]. Taken together, there is sufficient evidence to show variability between culture methods and donor influence hMSC behavior in 2D culture. However, there is a lack of this research in 3D biomaterials that include relevant extracellular matrix cues, and differences in hMSC response may be specific to the biomaterial being developed.

Our lab has developed an instructive 3D mineralized collagen scaffold that promotes osteogenesis *in vitro* and *in vivo* without the need for exogenous factors. We have shown these scaffolds promote osteogenesis *in vitro* using cells from animals [37–40] and humans [41–51], and improve bone healing *in vivo* using rat [52] and pig [53, 54] models. Further, our scaffolds have been used to answer other biological questions. Our collaborators have shown that MSCs cultured on the mineralized collagen scaffolds promote the secretion of osteoprotegerin, which subsequently reduces osteoclast resorption *in vitro* [49–51]. Moreover, our group has used these materials to understand the antimicrobial effects of manuka honey [55], the immunomodulatory effects of placenta-derived tissues [40, 44], and the crosstalk between immune cells and hMSCs [56]. Additionally, the material properties of these scaffolds are well studied and characterized in terms of stiffness (∼1000 kPa dry [38, 39], ∼30kPa, hydrated [45]), pore size (100-200 um [39, 40, 42]), and pore alignment (isotropic, anisotropic [42]). Given the comprehensive characterization of our mineralized collagen scaffolds and its demonstrated ability to induce osteogenesis, we believe our scaffold system is well suited to serve as a model material to study donor variability and its impact on osteogenic response *in vitro*.

In this manuscript, we compare differences in male and female hMSC activity and gene expression when cultured in a well-defined 3D mineralized collagen scaffold under development for craniofacial bone regeneration applications. We first examine the effect of cell passaging conditions (final passage number; total culture length between thawing from cryopreservation to scaffold seeding) for hMSCs from four donors (2 male, 2 female); we report changes in hMSC proliferation and secretion of pro-osteogenic factors during a short-term culture period (21 days). We subsequently examine the effect of donor and sex on proliferation, osteogenic gene expression, and soluble factor secretion using a larger group of hMSCs (4 male, 4 female) over a long-term culture experiment (56 days). To our knowledge this study is the first of its kind in evaluating the effect of passage scheme and donor sex and on hMSC osteogenic response in a 3D biomaterial.

## 2. Materials and Methods

### 2.1. Mineralized collagen scaffold fabrication

Mineralized collagen-glycosaminoglycan scaffolds were fabricated via lyophilization from a mineralized collagen precursor suspension. The mineralized collagen suspension was created by homogenizing type I collagen (Sigma Aldrich, St. Louis, Missouri USA), chondroitin-6-sulfate (Sigma Aldrich), and calcium salts (calcium hydroxide and calcium nitrate, Sigma Aldrich) in mineral buffer solution (0.1456M phosphoric acid/0.037M calcium hydroxide). The precursor suspension was stored at 4°C and degassed prior to lyophilization.

Scaffolds were subsequently fabricated via lyophilization using a Genesis freeze-dryer (VirTis, Gardener, New York USA) [57]. Briefly, 67.4mL of precursor suspension was pipetted into a custom stainless-steel mold (5 inches by 5 inches). Mineralized collagen sheets were then fabricated by freezing the suspension via cooling from 20°C to -10°C at a constant rate of 1°C/min followed by a temperature hold at -10°C for 2 hours. The frozen suspension was then sublimated at 0°C and 0.2 Torr, resulting in a porous scaffold network. After fabrication, a 6mm biopsy punch was used to obtain scaffolds that would be used for cell culture (6mm diameter and 3mm height). All scaffolds used for cell culture were sterilized via ethylene oxide treatment for 12 hours utilizing an AN74i Anprolene gas sterilizer (Andersen Sterilizers Inc., Haw River, North Carolina USA) in sterilization pouches [39, 45, 58]. All subsequent handling steps leading to studies of cell activity were performed in a sterile manner.

### 2.2. Mineralized collagen scaffold hydration

Dry sterile scaffolds were hydrated and crosslinked prior to cell seeding as previously described [37, 56, 59] and all steps were done under moderate shaking. Scaffolds were first hydrated for 2 hours in ethanol at ∼20°C (room temperature). Scaffolds were then washed in PBS for 1 hour at ∼20°C. Next, scaffolds were crosslinked for 2 hours in EDC-NHS at ∼20°C and washed again in PBS. Finally, the scaffolds were incubated in cell culture medium for 48 hours at 37°C with a complete media change after 24 hours.

### 2.3 Human mesenchymal stem cell culture

#### 2.3.1 Human mesenchymal stem cell (hMSC) source

Human mesenchymal stem cells (hMSCs) were purchased from Essent Biologics (Centennial, Colorado USA) and RoosterBio (Frederick, Maryland, USA). Cells from Essent Biologics are derived from cadaveric adipose tissue and cells from RooserBio are derived from bone marrow donors. **Table 1** contains donor ID, sex, age, ethnicity, pre-existing conditions, tissue origin, company, and lot number for each cell used. Passage 1 hMSCs were expanded in T175 flasks (Thermo Fisher Scientific, Hampton, New Hampshire USA) and were cultured until passage 3 and 4 in RoosterNourishTM-MSC expansion medium (RoosterBio) at 37°C and 5% CO2. Once at passage 3 and 4 cells were frozen and stored in liquid nitrogen until further use.

**Table 1:**
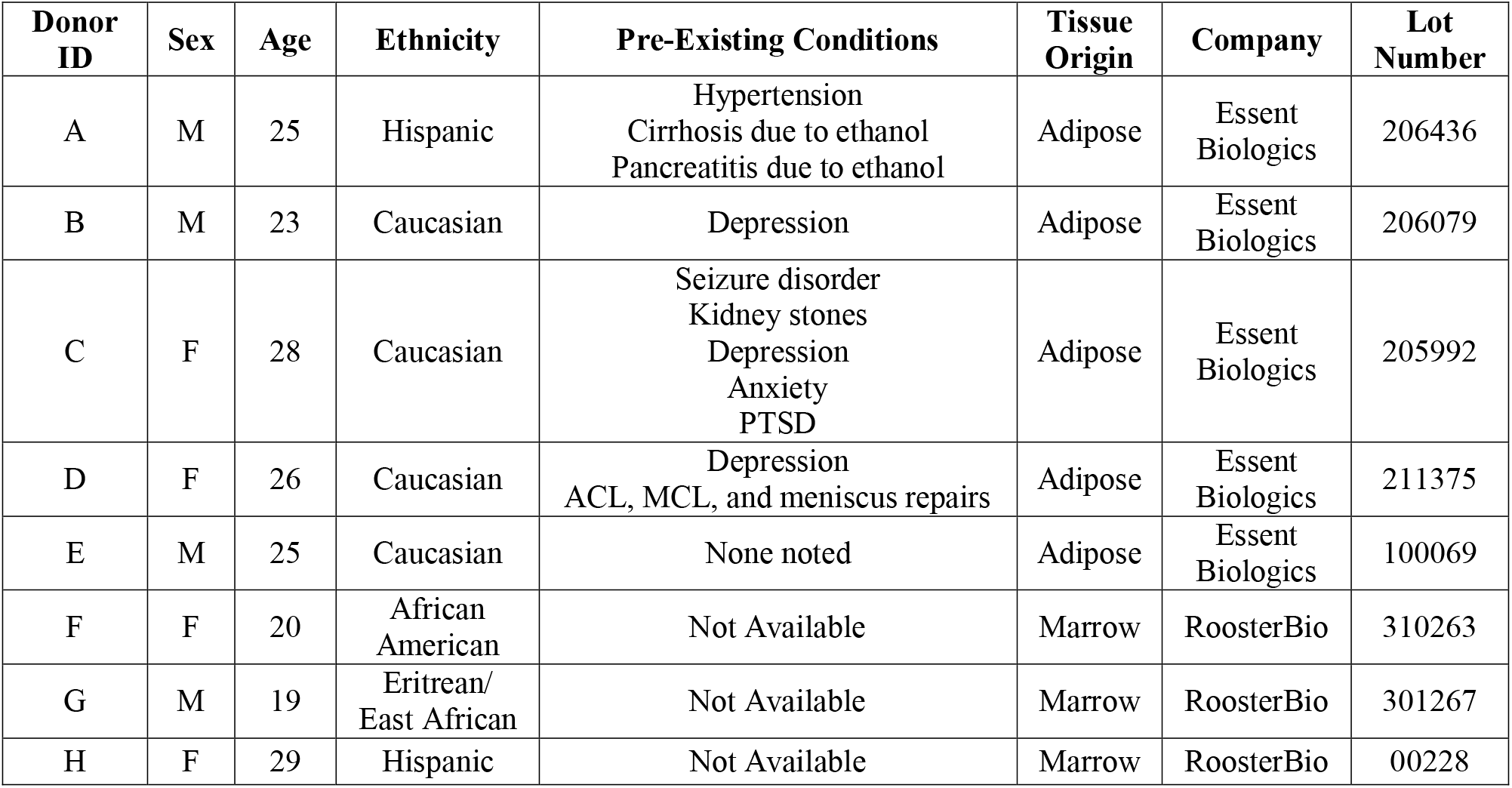
Donor information, company and tissue source, and lot numbers.

#### 2.3.2 Culturing hMSCs to compare passage number and culture time influences on cell response

Passage 3 hMSCs from 4 donors chosen randomly (2 male; B, G, and 2 female; C, D) were thawed and cultured in T175 flasks (Thermo Fisher Scientific, Hampton, New Hampshire USA) supplemented with RoosterNourishTM-MSC expansion medium (RoosterBio) at 37°C and 5% CO2. These passage 3 (p3) cells were expanded for two passages to p5 or for three passages to p6. These will be referenced as p3-5 and p3-6 for the rest of the study. Passage 4 hMSCs were thawed and cultured in T175 flasks supplemented with RoosterNourish^TM^-MSC expansion medium for one passage to p5. These cells with be referenced as p4-5 for the rest of the study. Once confluent, p3-5, p3-6, and p4-5 hMSCs were seeded (150,000 cells/scaffold) onto mineralized collagen scaffolds using an orbital seeding method [60, 61] for 6 hours at 37°C and 5% CO2. Cell seeded scaffolds were transferred to ultra-low attachment plates and cultured in mesenchymal stem cell growth media (phenol red free low glucose DMEM + L-glutamine, 10% fetal bovine serum, 1% antibiotic-antimycotic) at 37°C and 5% CO2 for 21 days (**Figure 1A**). Every three days, media was completely changed and 300uL of media was stored at -20°C for further analysis (ELISA).

**Figure 1:**
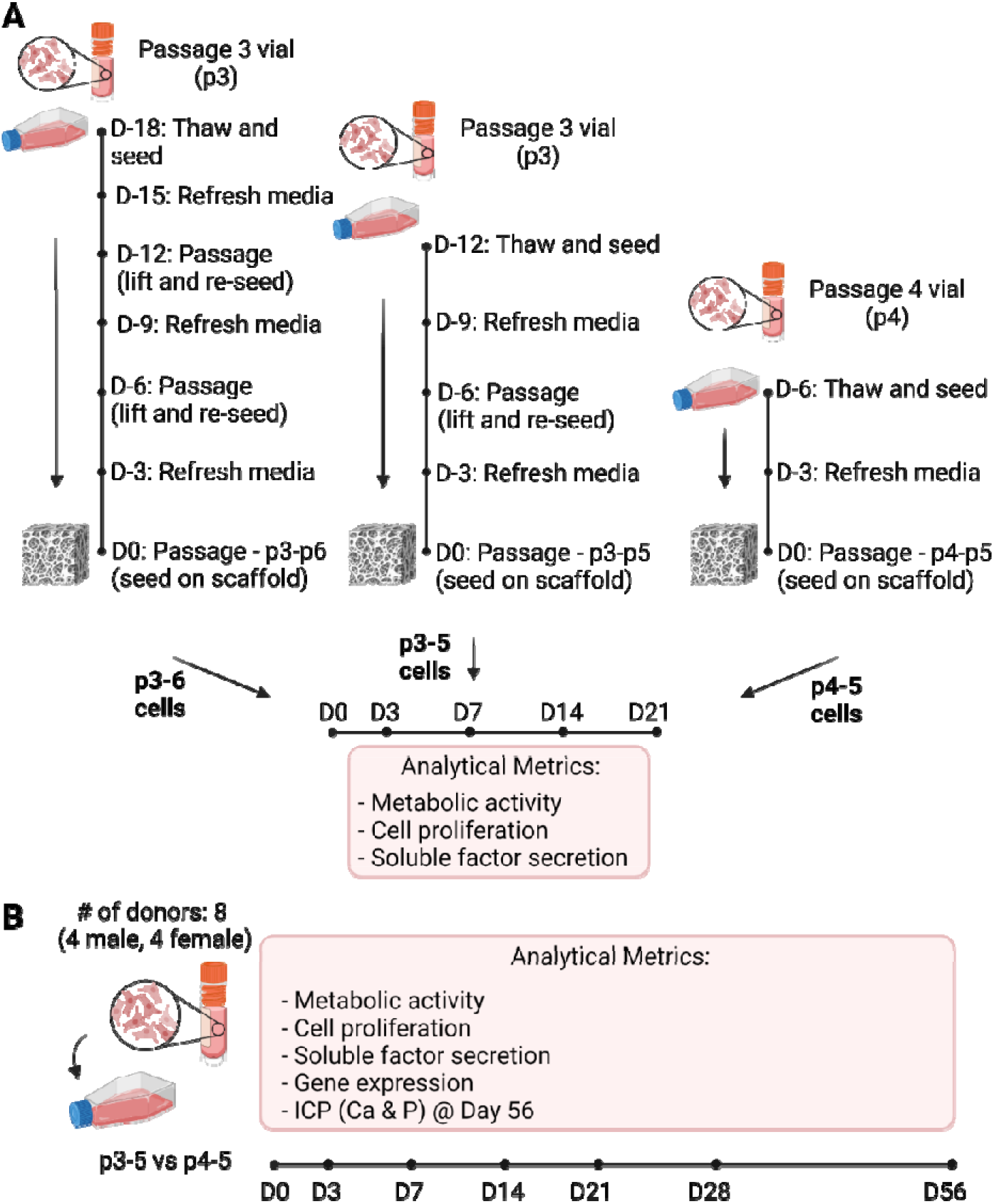
Experimental design and overview. (A) Experimental overview for passage scheme study. Human mesenchymal stem cells (hMSCs) from 4 donors (2 male, 2 female) were cultured on mineralized collagen scaffolds after varying passage schemes to compare the influence of passage number and culture length on cell proliferation and response. hMSCs were thawed at passage 3 and seeded at passage 6 (p3-6) undergoing 2 passages in culture prior to being seeded on mineralized collagen scaffolds. hMSCs were thawed at passage 3 and seeded at passage 5 (p3-5) undergoing 1 passage in culture prior to being seeded on mineralized collagen scaffolds. hMSCs were thawed at passage 4 and seeded at passage 5 (p4-5) undergoing 1 passage in culture prior to being seeded on mineralized collagen scaffolds. These cells were subsequently cultured for 21 days and their metabolic activity, cell proliferation and secretome were quantified. (B) Experimental overview for donor variability study. hMSCs from 8 donors (4 male, 4 female) were cultured as p3-5 or p4-5 groups and cultured on mineralized collagen scaffolds for 56 days. Cell response on these scaffolds was quantified to identify donor and sex-based differences.

#### 2.3.3 Culturing hMSCs to compare donor-donor variability

Passage 3 and passage 4 cells from 8 donors (4 male and 4 female) were expanded in T175 flasks (Thermo Fisher Scientific, Hampton, New Hampshire USA) and were cultured until passage 5 (p3-5 and p4-5, respectively) in RoosterNourish^TM^-MSC expansion medium (RoosterBio) at 37°C and 5% CO2. Once confluent, p3-5 and p4-5 hMSCs were seeded (150,000 cells/scaffold) onto mineralized collagen scaffolds using an orbital seeding method [60] for 6 hours at 37°C and 5% CO2. Cell seeded scaffolds were transferred to ultra-low attachment plates and cultured in mesenchymal stem cell growth media (phenol red free low glucose DMEM + L-glutamine, 10% fetal bovine serum, 1% antibiotic-antimycotic) at 37°C and 5% CO2 for 56 days (**Figure 1B**). Every three days, media was completely changed and 300uL of media was stored at -20°C for further analyses (ELISA, Luminex, ALP activity).

### 2.4 Cell Metabolic Activity & Cell Number

Metabolic activity was obtained using an alamarBlue viability assay (Invitrogen, Carlsbad, California USA) at Days 7, 14, and 21. Cell seeded scaffolds were incubated in a 10% alamarBlue solution for 90 minutes at 37°C under moderate shaking. The fluorescence of the alamarBlue solution was then read using a fluorescent spectrophotometer (Tecan Infinite F200 Pro, Männedorf, Switzerland). The active ingredient in the alamarBlue reagent, resazurin, is reduced to a compound that is highly fluorescent when added to metabolically active cells [45, 62]. The relative cell metabolic activity was determined from standard curves generated with known hMSC concentrations for p3-5, p3-6, and p4-5. The relative cell metabolic activity was reported as a fraction of initial seeding cell count (i.e., an experimental value of 1 indicates the metabolic activity of the number of cells seeded onto the scaffold).

Cell numbers were obtained via DNA isolation at Days 7, 14, and 21. DNA extraction was completed using a DNeasy Blood & Tissue Kit (Qiagen, Hilden, Germany). Cell-seeded scaffolds were cut into quarters and placed into 360uL Buffer ATL and 40uL proteinase K and incubated at 56°C for 18-24 hours. Then, 400uL Buffer AL and 400uL ethanol were added to the lysates and vortexed. Following steps were followed per manufactures instructions. DNA concentration was quantified using a NanoDrop Lite Spectrophotometer (Thermo Fisher Scientific) and cell number was determined from standard curves generated with known cell numbers for p3-5, p3-6, and p4-5.

### 2.5 RNA isolation from mineralized collagen scaffolds

RNA was isolated from cells at Day 0 using the RNAqueous™ Total RNA Isolation Kit (Invitrogen), and the RNA was eluted in 40uL Elution Solution. RNA was isolated from cell-seeded scaffolds using TRIzol (Thermo Fisher Scientific) and the RNeasy Mini Kit (Qiagen). Cell-seeded scaffolds were cut into quarters and placed into Phasemaker™ Tubes (Invitrogen). Next, 1mL TRIzol was added to each tube, scaffolds with TRIzol were vortexed and incubated at room temperature for 5 minutes. Then, 200uL chloroform was added to each tube, and scaffolds with TRIzol and chloroform were vortexed and incubated at room temperature for 3 minutes. Tubes were vortexed immediately before centrifuging at 15,000g for 15 minutes at 4LC. After centrifugation, the aqueous phase was added to 650uL 70% ethanol and mixed briefly. This solution was then pipetted into the RNeasy Mini Kit extraction columns and the kit instructions were followed as per manufacturers recommendation. RNA was eluted in RNAse free water and quantified using a NanoDrop Lite Spectrophotometer (Thermo Fisher Scientific). RNA samples were stored at -80LC until further analysis.

### 2.6 Nanostring gene expression evaluation

Transcript expression was quantified with the NanoString nCounter System (NanoString Technologies, Inc.) located at the Tumor Engineering and Phenotyping Shared Resource at the Cancer Center at Illinois using a custom panel of 38 mRNA probes (**Supplemental Table 1**). The NanoString nCounter System identifies and counts individual transcripts using unique color-coded probes, and there is no reverse transcription or amplification steps required. Isolated RNA was quantified and quality tested using Qubit RNA BR Assay Kit, loaded into cartridges, and run on the NanoString assay as instructed by the manufacturer. The nSolver Analysis Software (NanoString Technologies, Inc.) was used for data processing, normalization, and evaluation of expression. Raw data was normalized to Day 0 controls (n=3). Expression levels are depicted as log2 fold change.

### 2.7 Enzyme-linked immunosorbent and Luminex assays

An enzyme-linked immunosorbent assay (ELISA) (R&D Systems, Minneapolis, MN) was used to quantify the amount of osteoprotegerin (OPG) released by cells seeded on mineralized collagen scaffolds. Media was pooled into the following groups: Day 3, Days 6-9, Days 10-15, Days 16-30, Days 33-56. Similarly, a custom Luminex panel (R&D Systems) was used to quantify the amount of 14 proteins released by cells seeded on mineralized collagen scaffolds (**Supplemental Table 2)**. Media was pooled as described above for the ELISA (n=4). The assay was conducted as per manufacturers protocol and absorbance was read on a Luminex 200 XMAP system (Luminex Corporation, Austin, TX) and each analyte was normalized to a background control and converted to a concentration using its own standard curve. Expression levels of the analytes are depicted as cumulative concentrations.

### 2.8 Alkaline phosphatase (ALP) Assay

An ALP activity assay (Abcam, England) was used to determine cell-dependent osteogenic activity (n=6). Results were compared between media isolated from cell-seeded scaffold groups for the entire length of culture, using unseeded scaffold groups as a background control. P-nitrophenyl phosphate (pNPP, µmol) concentration per well was converted to U/well with known reaction time and volume of sample. Media was collected every 3 days and pooled into the following groups for analysis: Day 3, Days 6-9, Days 10-15, Days 16-30, Days 33-56.

### 2.9 Quantification of Calcium and Phosphorus in freeze-dried scaffolds

Scaffolds for each donor were harvested on Day 56 (n=6), washed in PBS, fixed in formal-fix at 4°C overnight, and washed in PBS three times for 5 minutes at room temperature under moderate shaking. The scaffolds were then frozen at -80°C overnight and freeze-dried using a Genesis freeze-dryer (VirTis, Gardener, New York USA). Freeze-dried scaffold specimens were analyzed via inductively coupled plasma optical emission spectrometry (ICP-OES). Dry samples were transferred to a digestion tube, then digested with concentrated nitric acid (67-70%) followed by automated sequential microwave digestion in a CEM Mars 6 microwave digester (CEM Microwave Technology Ltd., North Carolina, USA). The final product was a clear aqueous digest which was diluted to a volume of 25mL using DI water. This solution was then introduced to inductively coupled plasma-mass spectrometer (NexION™ 350D ICP-MS, PerkinElmer, USA) for the elemental analysis in a standard mode.

### 2.10 Statistics

RStudio was used for all plotting (ggplot2) and statistical analysis. No outliers were removed. Sample size was six (n=6) for alamarBlue, ELISA, ALP, and ICP-OES, four (n=4) for Luminex, and three (n=3) for NanoString and cell number. All experiments had three or more experimental groups. Residuals were tested for normality using the Shapiro-Wilk test, and homogeneity of variance was tested using the Levene Test.

For passage scheme experiments (**Figure 2**): when both assumptions were met we ran a one-way ANOVA (independent variable was passage scheme) and Tukey’s HSD mean separation with alpha=0.05, when data did not have equal variance we ran a one-way ANOVA with Welch’s correction and Tukey’s HSD mean separation with alpha=0.05, when data were not normally distributed we ran a one-way ANOVA and Tukey’s HSD mean separation with alpha=0.01, and when both assumptions were unmet we ran Mood’s median for non-parametric data and median separation with alpha=0.05. For donor variability experiments (**Figures 3-7**): when both assumptions were met we ran a two-way ANOVA (independent variables were donor and sex) and Tukey’s HSD mean separation with alpha=0.05, and if one or both assumptions were unmet we ran a two-way ANOVA and Tukey’s HSD mean separation with alpha=0.01.

**Figure 2:**
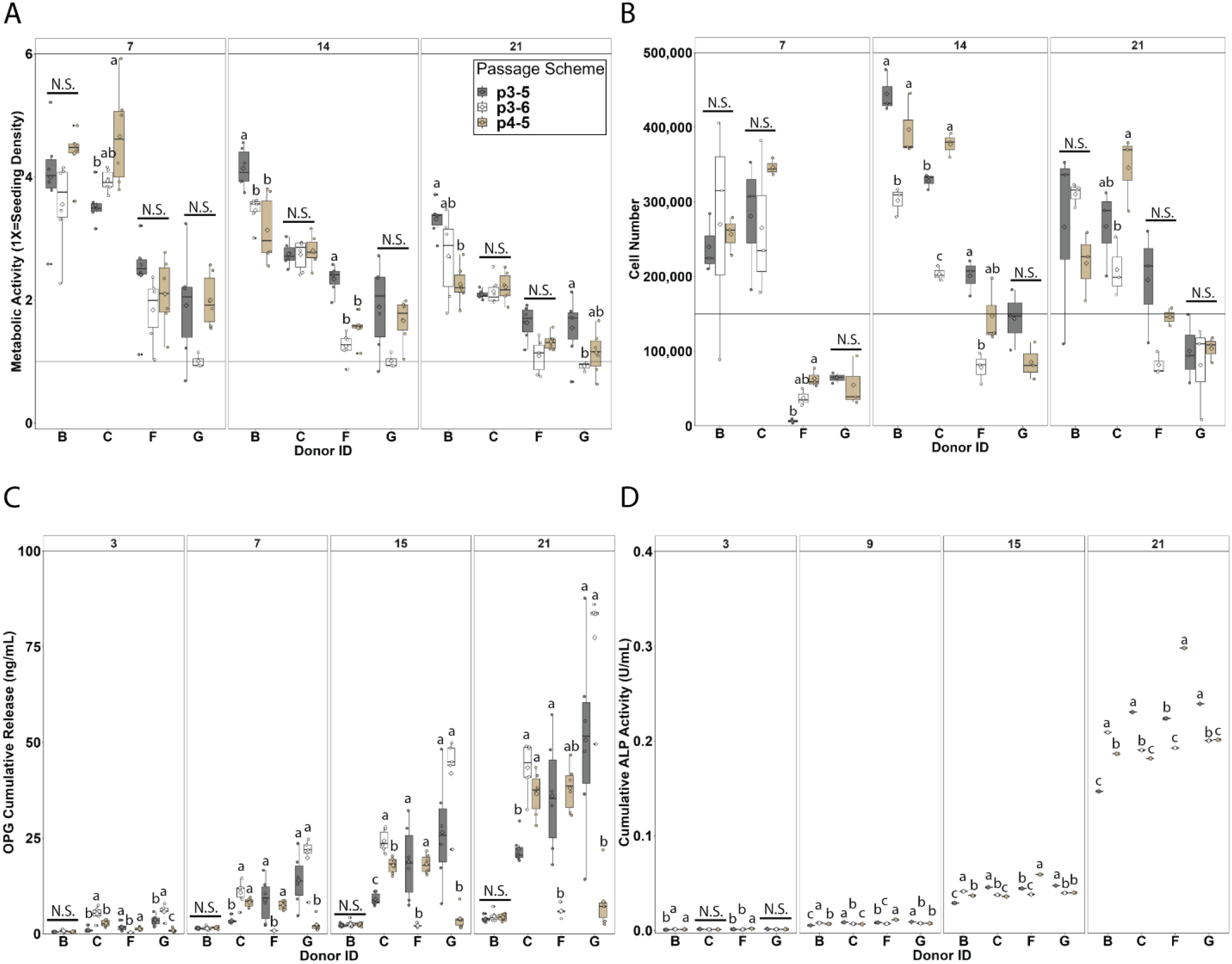
Passage scheme influences on metabolic activity, cell number, soluble factor secretion, and ALP activity. Four donors (B, G – male, and C, F – female) with three passage methods (p3-5, p3-6, and p4-5) were cultured on mineralized collagen scaffolds for 21-days. (A) Metabolic activity was quantified via alamarBlue and decreases over time for all donors. The effect of passage scheme on metabolic activity is donor dependent. (B) Cell number was quantified via DNA extraction and remains relatively constant for all donors. The effect of passage scheme on cell number is donor dependent. (C) Secreted osteoprotegerin (OPG) was quantified using an ELISA and increases over 21 days in culture for all donors. Secretion is significantly different between passages at all timepoints for all donors except Donor B. (D) Alkaline phosphatase (ALP) activity was quantified using an ALP activity kit and increases over the 21 days of culture for all donors. ALP activity is significantly different for all donors beginning at Day 9. Groups that share a letter are not significantly different (p<0.05). Alphabetical order indicates magnitude (i.e., group labeled a has the highest experimental value).

**Figure 3:**
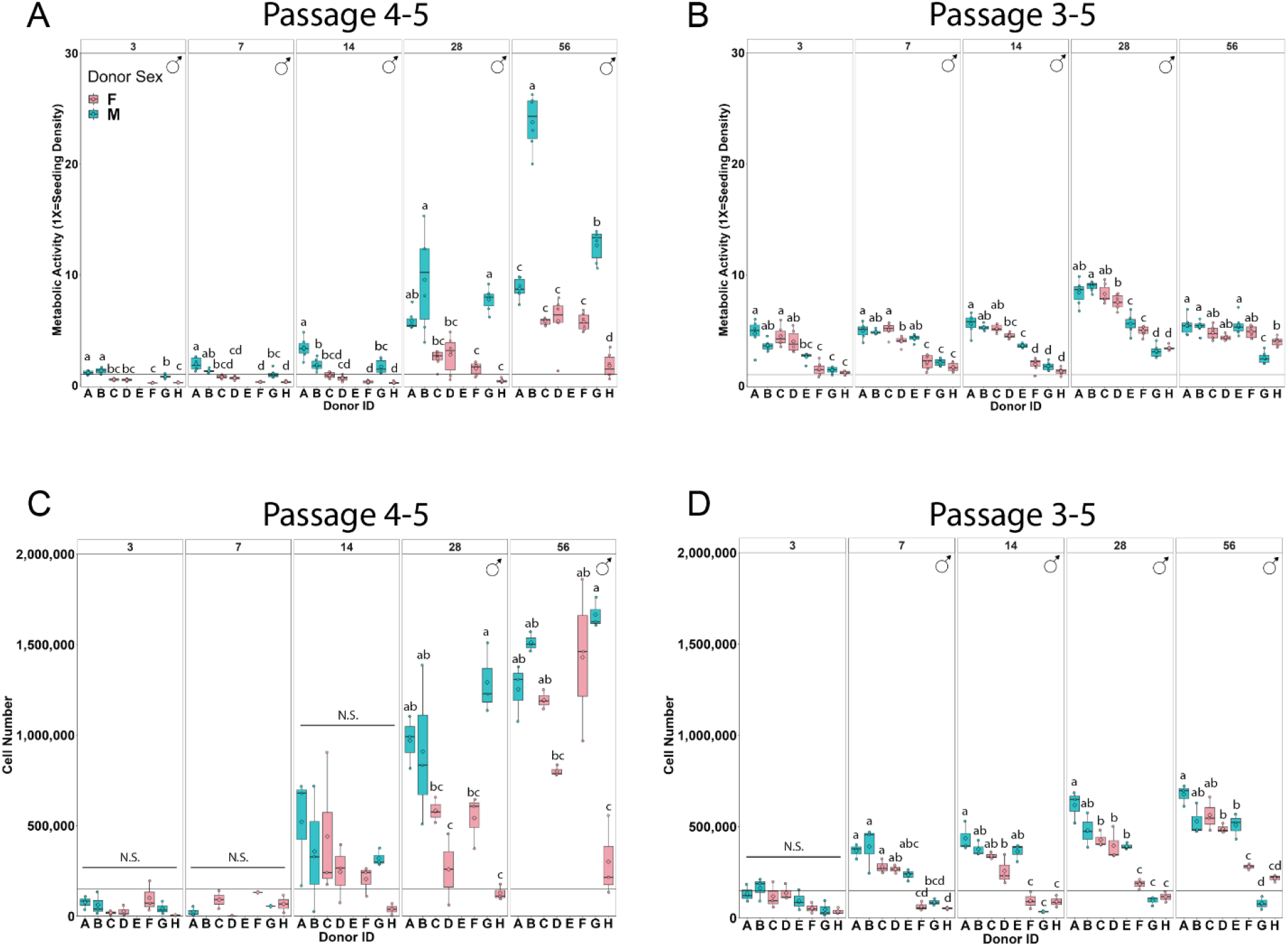
Donor variability influence on metabolic activity and cell number. Eight donors (A, B, E, G – male, and C, D, F, H – female) with 2 passage methods (p4-5 and p3-5) were cultured on mineralized collagen scaffolds for 56-days. Metabolic activity and cell number were quantified over the course of the experiment. (A, B) Metabolic activity was quantified via alamarBlue. Male cells are more metabolically active than female cells in both passage schemes. (C, D) Cell number was quantified via DNA extraction. Male cells are more proliferative than female cells in both passage schemes. Groups that share a letter are not significantly different (p<0.05). Alphabetical order indicates magnitude (i.e., group labeled a has the highest experimental value). Sex symbol (male: ♂, female: ♀) indicates sex-based significance (p<0.05) at the indicated timepoint.

## 3. Results

### 3.1. Passage scheme influence on metabolic activity and cell number is donor dependent

Metabolic activity and cell number were quantified for 21 days of culture (**Figure 2**). The impact of passage scheme on metabolic activity is donor dependent while optimal passage scheme varied between donors and appeared to be inconsistent within sex. All 4 donors (B: male, C: female, G: male, and F: female) and passage schemes (p3-5: thawed at passage 3 and seeded at passage 5, p3-6: thawed at passage 3 and seeded at passage 6, and p4-5: thawed at passage 4 and seeded at passage 5) displayed decreasing metabolic activity with time (**Figure 2A**). The most dramatic decrease from Day 7 to Day 21 was observed for donors B (male) and C (female); however, these donors (B, C) displayed the highest metabolic activity at each timepoint. Although no trend as a function of passage scheme was observed across all donors, metabolic activity varied as a function of passage scheme for all donors at varying time points. Significant differences in metabolic activity between passage schemes were observed at Day 7 for donor C (female, p=0.02124), at Day 14 for donors B (male, p=0.00142) and F (female, p=1.3e-6), and at Day 21 for donors B (male, p=0.00381) and G (male, p=0.00499).

Cell number for each donor remained relatively constant with time except for donor F (female) which displayed increasing cell number (**Figure 2B**). Donors B (male) and C (female) displayed the highest cell number which mirrors their higher metabolic activity. There was no trend as a function of passage scheme observed across all donors, and cell number varied as a function of passage scheme for donors B, F, and G at varying time points. Significant differences in cell number between passages were observed at Day 7 for Donor F (female, p= 0.00645), at Day 14 for donors B (male, p=0.00387), C (female, p=7.25e-06), and F (female, p=0.00882), and at Day 21 for donor C (female, p=0.0427). Passage scheme p3-6 appeared to be the worst for cell number in all donors, resulting in too few cells for seeding donor G (male) for days 7 and 14 cell number analysis and having the lowest cell number for donors C (female) and F (female) at days 14 and 21 and donor B (male) at day 14. To summarize, these data show that metabolic activity and cell number do not correlate across all passage schemes and donors.

### 3.2. Passage scheme significantly impacts hMSC osteogenic response

Osteoprotegerin (OPG) and alkaline phosphatase activity (ALP) were quantified in their soluble form over the 21-day culture period (**Figure 2**). OPG is a soluble decoy receptor to RANKL inhibiting osteoclastogenesis and bone resorption, and the MC scaffolds used for these studies have previously been shown to promote endogenous production of OPG by MSCs. Passage scheme significantly impacts OPG secretion. OPG secretion increased with time across all donors with the least dramatic increase in donor B (male) (**Figure 2C**). Passage scheme has a significant impact on OPG secretion at all time points for donors C (male, Day 3 p=0.000106 | Day 7 p=4.49e-05 | Day 15 p= 2.62e-08 | Day 21 p=2.72e-05), F (female, Day 3 p=0.009404 | Day 7 p=0.009404| Day 15 p= 0.009404 | Day 21 p=1.297e-05), and G (male, Day 3 p=0.000143 | Day 7 p=0.000356 | Day 15 p=0.000118 | Day 21 p=4.77e-05). Although no trend in passage scheme was observed, passage scheme p3-5 displayed the most variable OPG secretion for donors G (male) and F (female), with donor G displaying the greatest variability in OPG secretion across passage schemes.

ALP activity increased with time for all donors and passage scheme significantly impacts ALP activity (**Figure 2D**). All donors displayed similar amounts of ALP expression across timepoints despite differences in their metabolic activity and cell number. Passage scheme significantly impacted ALP activity at Day 3 for donors B (male, p=0.00278) and F (female, p=0.000963), at Day 9 for donors B (male, p=2.02e-12), C (female, p=6.74e-11), F (female, p=7.41e-15), and G (male, p=1.66e-08), at Day 15 for Donors B (male), C (female), F (female), and G (male) (p<2e-16 for all donors), and at Day 21 for donors B (male), C (female), F (female), and G (male) (p<2e-16 for all donors).

Overall, passage scheme appears to significantly influence cell activity and soluble factor secretion, but minimally impacts ALP activity. Taken together, no one passage scheme was optimal for enhanced metabolic activity, proliferation, and osteogenic factor secretion but rather these metrics were donor dependent. For the next step of this work, we evaluated donor variability and chose to move forward with passage schemes p4-5 and p3-5 only. This is based on passage scheme p3-6 having too few cells for cell seeding in Donor G and being consistently low for metabolic activity (**Figure 2A**) and cell number (**Figure 2B**).

### 3.3. Metabolic activity and cell number vary between donors and display significant sex differences

Metabolic activity and cell number were quantified for all eight donors over the course of 56 days in both p4-5 (thawed at passage 4 and seeded at passage 5) and p3-5 (thawed at passage 3 and seeded at passage 5) passage schemes (**Figure 3**). All donors displayed increasing metabolic activity with time with a more dramatic increase in p4-5 groups (**Figure 3A,B**). Cells from male donors displayed significantly higher metabolic activity than cells from female donors at days 3 (p< 2e-16), 7 (p= 5.71e-12), 14 (p= 2.24e-13), 28 (p= 1.26e-11), and 56 (p <2e-16) in passage scheme p4-5 and at days 7 (p=8.4e-07), 14 (p=2.29e-07), and 28 (p=0.0257) in passage scheme p3-5. Metabolic activity varied between donors at all timepoints in both passage scheme p4-5 (Day 3 p=8.41e-07, Day 7 p=2.24e-05, Day 14 p=6.83e-06, Day 28 p=0.0159, Day 56 p<2e-16) and p3-5 (Day 3 p=1.5e-12, Day 7 p< 2e-16, Day 14 p< 2e-16, Day 28 p< 2e-16, day 56 p=8.14e-10) with variability between donors of the same sex; thus, while this cohort of male cells was significantly more metabolically active than this cohort of female cells, there is still variability between donors of the same sex. Passage scheme p4-5 had higher and more variable metabolic activity than p3-5 with donor B (male) displaying significantly higher metabolic activity compared to all other donors at Day 56 in p4-5, surpassing a 20-fold increase compared to Day 0. Furthermore, donor H (female) consistently displayed the lowest metabolic activity and cell number throughout the cell culture in p4-5. These data show that sex differences are more apparent in the p4-5 group with greater metabolic activity changes; however, sex differences still reveal themselves in the p3-5 group.

All donors displayed increasing cell number with time with a more dramatic increase in passage scheme p4-5 (**Figure 3C,D**). Male donors displayed higher cell number than female donors at days 28 (p=4.78e-06) and 56 (p=2.81e-05) in passage scheme p4-5 and at days 7 (p=0.000155), 14 (p=9.58e-05), 28 (p=0.000278), and 56 (p=0.0203) in passage scheme p3-5. Cell number varied significantly between donors at days 28 (p=0.0481) and 56 (p=0.000123) in passage scheme p4-5 and at days 7 (p=1.46e-06), 14 (p=1.31e-07), 28 (p=8.49e-08), and 56 (p=7.08e-09) in passage scheme p3-5 with variability between donors of the same sex. Like the metabolic activity data, while this cohort of male cells had significantly more cells than this cohort of female cells, there is still variability between donors of the same sex. Passage scheme p4-5 had higher and more variable cell number than p3-5, and this mirrors the higher and more variable metabolic activity observed in p4-5 compared to p3-5. To summarize, these data show that male donors tend to display greater metabolic activity and cell proliferation compared to female donors, however, these metrics do not correlate to each other, and relative values vary widely across passage scheme.

### 3.4. Donor and sex-based significance osteoprotegerin secretion and alkaline phosphatase activity is dependent on passage scheme

Osteoprotegerin (OPG) secretion and alkaline phosphatase (ALP) activity were quantified throughout the 56-day experiment for all eight donors for both p4-5 and p3-5 groups. All donors displayed increasing OPG secretion with time with a more dramatic increase in passage 4-5 (**Figure 4A, B**). Male cells had significantly higher OPG secretion than female cells at Day 3 in passage scheme p4-5 (p= 0.00719), but no sex differences were observed in the following timepoints, and no sex differences were observed in passage scheme p3-5. Significant donor variability was observed in passage scheme p4-5 at all timepoints (Day 3 p=1.94e-08, Day 7 p=5.38e-10, Day 14 p=2.84e-12, Day 28 p<2e-16, Day 56 p=7.05e-15) but not in p3-5 aligning with trends observed in the passage scheme experiment (**Figure 2**).

**Figure 4:**
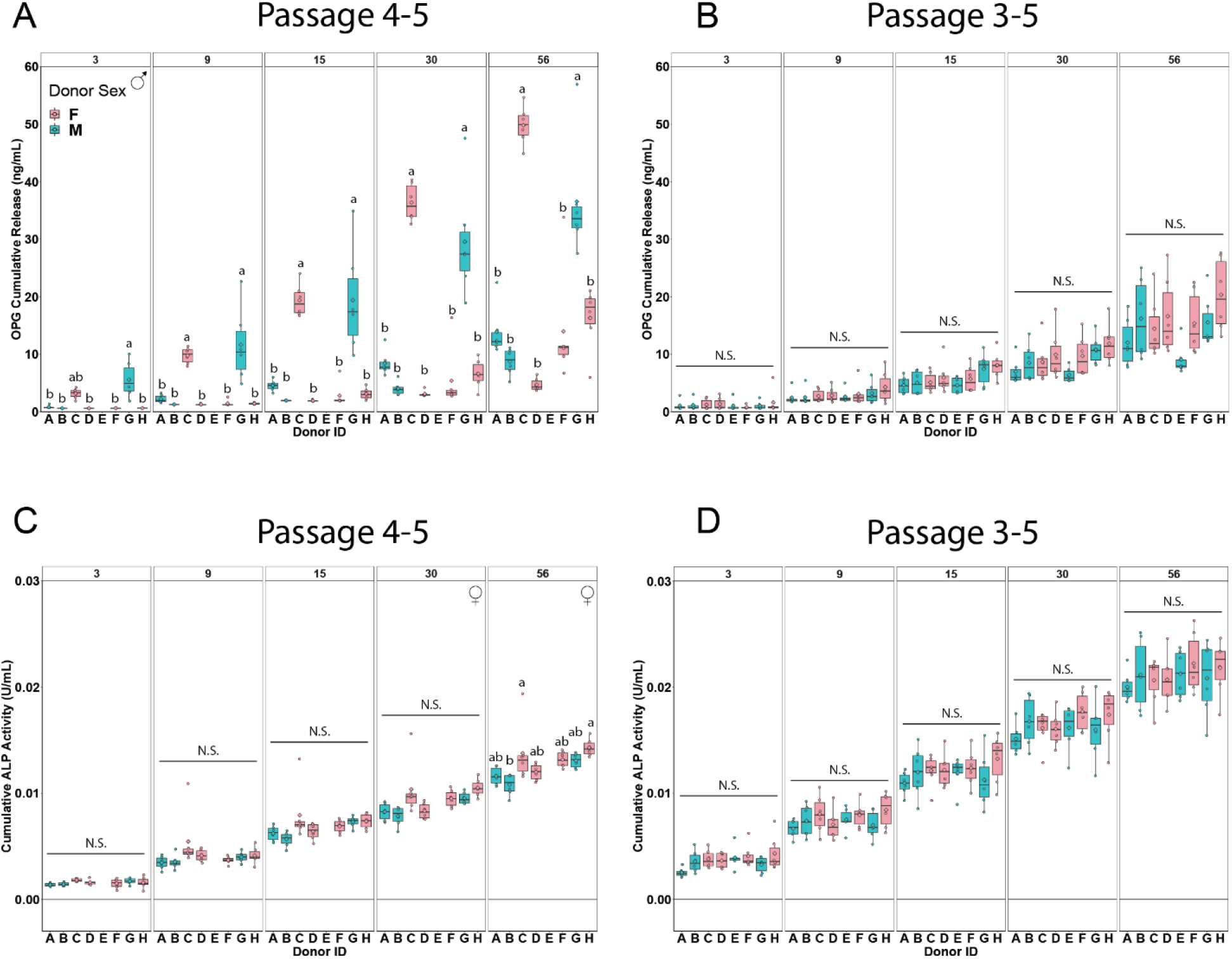
Donor variability influence on osteoprotegerin secretion and alkaline phosphatase activity. Eight donors (A, B, E, G – male, and C, D, F, H – female) with 2 passage methods (p4-5 and p3-5) were cultured on mineralized collagen scaffolds for 56-days. (A, B) Osteoprotegerin (OPG) was quantified via ELISA. There were no significant sex differences in OPG secretion for either passage scheme. (C, D) Alkaline phosphatase (ALP) activity was quantified using an ALP activity assay. Female cells have significantly higher ALP activity than male cells in p4-5. Groups that share a letter are not significantly different (p<0.05). Alphabetical order indicates magnitude (i.e., group labeled a has the highest experimental value). Sex symbol (male: ♂, female: ♀) indicates sex-based significance (p<0.05) at the indicated timepoint.

All donors displayed increasing ALP activity with time with activity appearing similar between passage schemes (**Figure 4C, D**). Sex differences were observed in passage scheme p4-5 at days 28 (p=0.00570) and 56 (p=0.00103) with female cells displaying higher ALP activity compared to male cells (**Figure 4C**). However, no impact of sex on ALP activity was observed in passage scheme p3-5 (**Figure 4D**). Once again, sex differences being present in passage scheme p4-5 but not p3-5 remains consistent when evaluating osteogenic output. In p4-5 donor variability was present only at Day 56 (p= 0.00694), and there was no significance between donors in p3-5. To summarize, these data show that passage scheme has a significant impact on osteogenic functional assays.

### 3.5. Female donors display significantly more mineral deposition compared to male donors

Mineral remodeling of the cell-seeded scaffolds was determined as the weight percent of calcium and phosphorous in the cell-seeded mineralized collagen scaffolds measured at Day 56 and compared to an unseeded Day 56 control. While a proxies for measuring changes in calcium phosphate content over the course of the experiment, this approach provides a direct measurement of osteogenic activity within the scaffold. Scaffolds seeded with female cells displayed significantly more calcium compared to those seeded with male cells in passage scheme p4-5 (p=2.3e-14) and significantly more calcium than the unseeded control in both passage schemes (**Figure 5A, B)**. Donor H (female), which notably displayed the lowest cell number and metabolic activity, displayed the highest calcium content across all donors in passage scheme p4-5 d. Scaffolds seeded with female cells also displayed significantly more phosphorus compared to scaffold seeded with male cells in passage scheme p4-5 (p=0.00325), with donor H (female) again displaying the highest phosphorous content. All donors have less phosphorus compared to the unseeded control in passage scheme p3-5 (**Figure 5C,D**). To summarize, these data show that cells from female donors display a strong osteogenic response (ALP activity, mineral content) in our mineralized collagen scaffolds despite having lower cell activity and proliferation than cells from male donors. This is apparent in both passage schemes given cell metabolic activity and cell number were significantly higher in this cohort of male cells (**Figure 3**), but there were no sex-based differences in ALP activity (**Figure 4**) and higher (p4-5) or equal (p3-5) amounts of mineral deposition (**Figure 4**) in this cohort of female cells.

**Figure 5:**
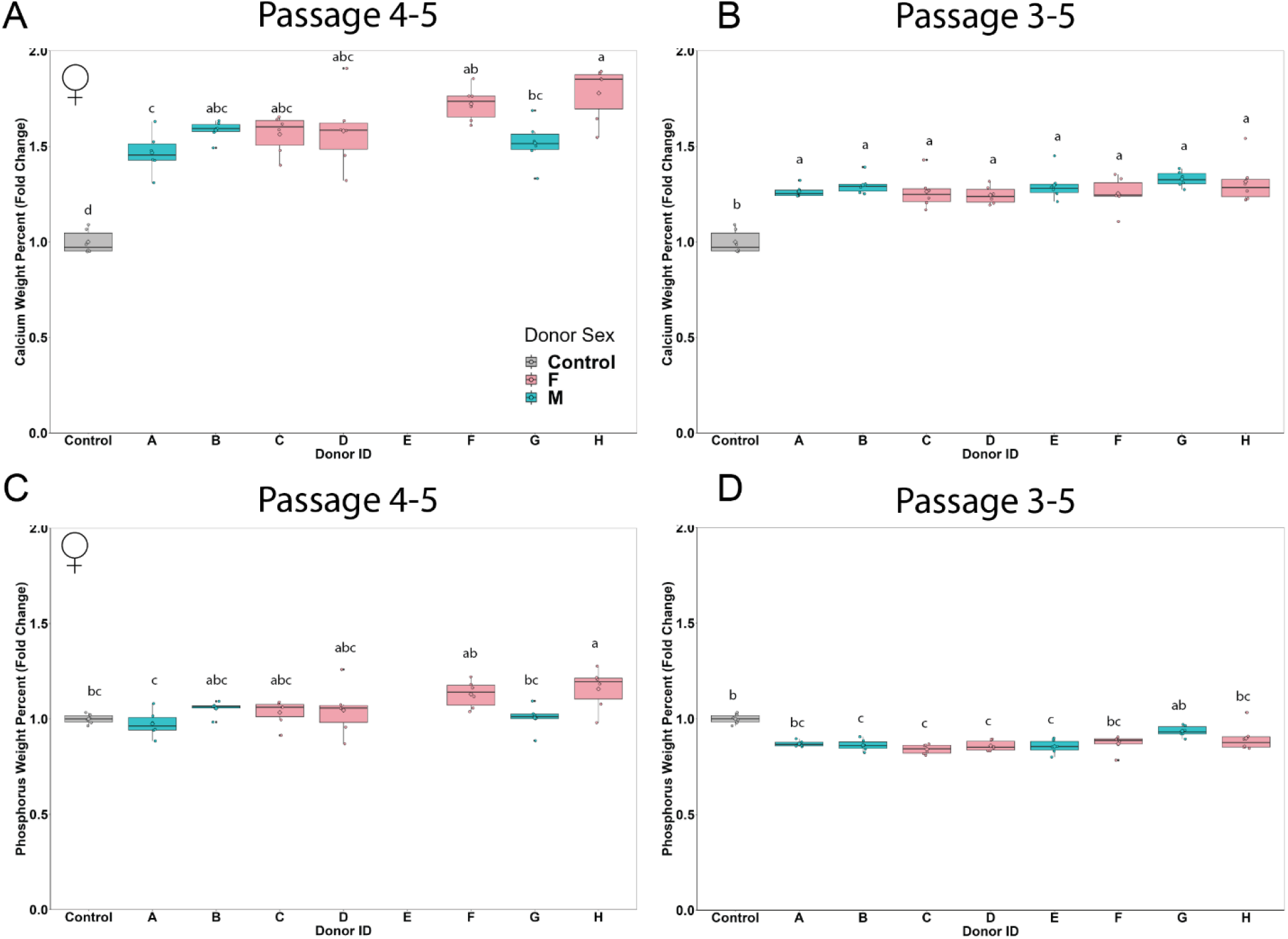
Donor variability influence on calcium and phosphorus content in mineralized collagen scaffolds. Eight donors (A, B, E, G – male, and C, D, F, H – female) with 2 passage methods (p4-5 and p3-5) were cultured on mineralized collagen scaffolds for 56-days. Mineral content was quantified at Day 56. Cell-seeded scaffolds are compared to unseeded control scaffolds that underwent all media changes throughout the 56-day culture period. (A, B) Calcium content at Day 56 was quantified using inductively couple plasma mass spectrometry (ICP-MS). Female cells have higher calcium content than male cells in p4-5. (C, D) Phosphorous content was quantified using ICP-MS. Female cells have higher phosphorus content than male ells in p4-5. Groups that share a letter are not significantly different (p<0.05). Alphabetical order indicates magnitude (i.e., group labeled a has the highest experimental value). Sex symbol (male: ♂, female: ♀) indicates sex-based significance (p<0.05) at the indicated timepoint.

### 3.6. Male donors display significantly higher osteogenic and immunomodulatory gene expression compared to female donors

We subsequently quantified expression patterns for osteogenic, immunomodulatory, and angiogenic genes for all 8 donors throughout the 56-day culture period for p4-5 and p3-5 groups using a custom 38 gene NanoString panel (35 functional genes and 3 housekeeping genes). Here, we have plotted gene expression at Day 56 (**Figure 6A,C,E**). We also took the difference in fold-change between passage schemes (absolute value of p4-5 fold-change subtracted from p3-5 fold-change) and represented those delta values in a heat map (**Figure 6B,D,F**). Please see **Supplemental Figures 1-3** for gene expression data across all timepoints and **Supplemental Table 3** for p-values and sex-based significance for all NanoString data.

**Figure 6:**
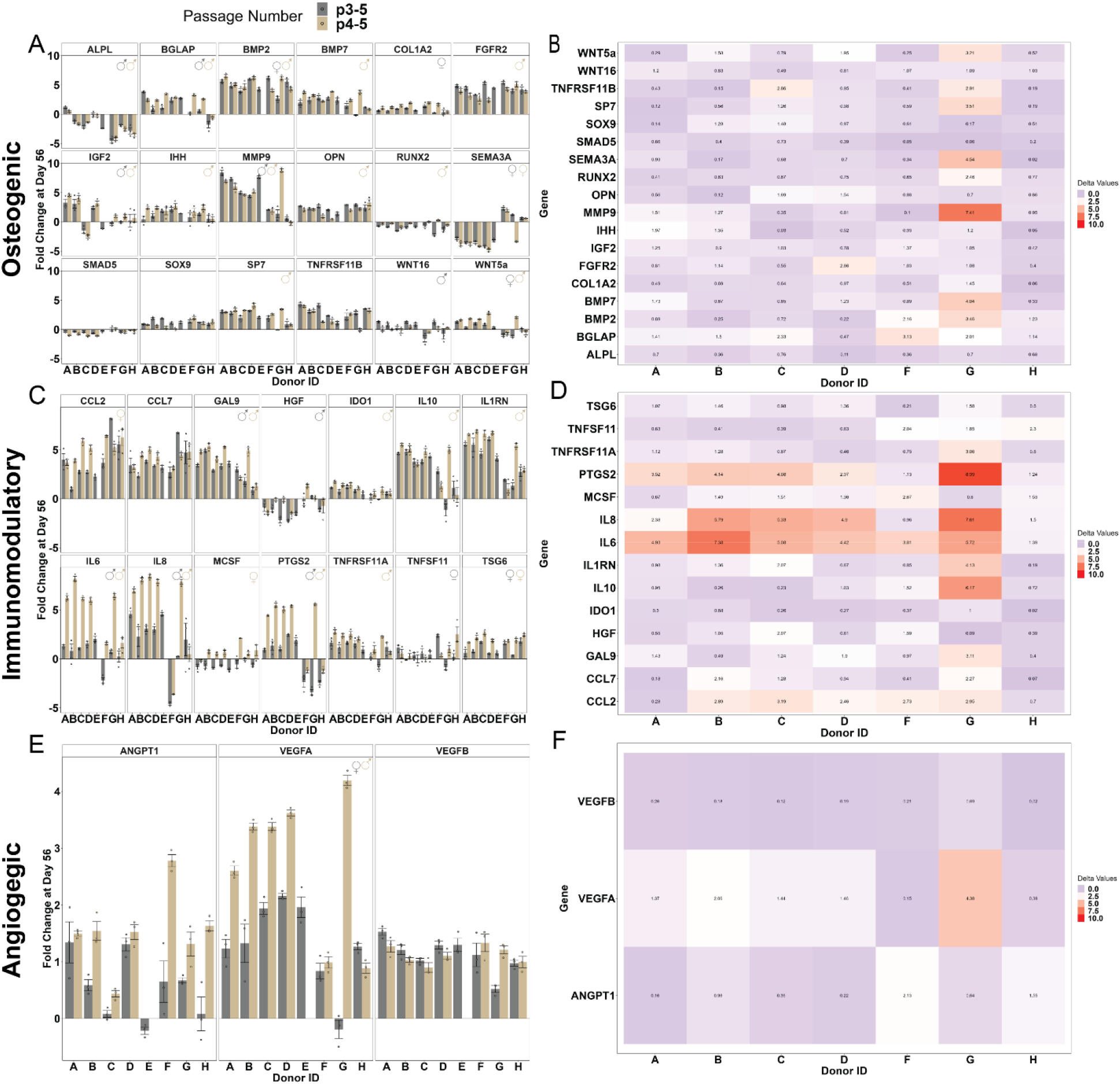
Gene expression analysis at Day 56 of culture. Eight donors (A, B, E, G – male, and C, D, F, H – female) with 2 passage methods (p4-5 and p3-5) were cultured on mineralized collagen scaffolds for 56-days. A custom Nanostring gene expression panel was used to quantify gene expression over the course of the experiment. (A) Male cells have higher expression of osteogenic genes in both passage schemes, (C) Male cells express significantly higher amounts of immunomodulatory genes in both passage schemes, (E) Male cells express significantly higher amounts of angiogenic genes in both passage schemes. Heatmaps were generated to visualize differences in fold change between the p4-5 and p3-5 schemes. (B) Osteogenic genes varied the most between passage schemes for Donors E and G, (D) Immunomodulatory genes varied between most donors with interleukin-6 and -8 being the most variable across donors. (F) Angiogenic gene expression had little variability between passage schemes. Sex symbol (male: ♂, female: ♀) indicates sex-based significance (p<0.05) at the indicated timepoint. Fold change is plotted as the mean +/- standard error.

First, we observe that male cells display higher osteogenic gene expression compared to female cells in both passage schemes at Day 56 (**Figure 6A**). Specifically, 12 out of 18 osteogenic genes in p4-5 and 5 out of 18 genes in p3-5 were expressed significantly higher in male cells. These data further indicate that the p4-5 passage scheme displays more frequent sex-based differences than the p3-5 passage scheme. Donors E and G displayed the greatest differences in fold change between passage schemes (**Figure 6B**). Here, we observe the difference in fold-change between passage schemes is less than 2 for most osteogenic genes evaluated (**Figure 6B**). Further, 117 out of 126 (93%) of osteogenic fold-change comparisons between p4-5 and p3-5 differ by less than one order of magnitude (**Supplemental Table 4**) and 12 out of 18 (67%) osteogenic genes have low variability between passage schemes with 3 or fewer donors (less than 50%) showing significant differences in fold-change (**Supplemental Table 5**). Notably, BMP2 showed reversed sex-based differences based on passage scheme with female cells expressing significantly higher levels in the p3-5 scheme while male cells expressed significantly higher levels in the p4-5 scheme.

Male cells also displayed generally higher immunomodulatory gene expression than female cells at Day 56 (**Figure 6C**). Specifically, 8 out of 14 immunomodulatory genes in p4-5 and 5 out of 14 genes in p3-5 were significantly higher in male cells. Again, these data indicate that the p4-5 passage scheme displays greater sex-based differences than the p3-5 passage scheme (with donors B, C, D, E, and G displayed the greatest differences in fold change between passage schemes; **Figure 6D**). The difference in gene expression fold change between passage schemes was less than 2 for more than half of immunomodulatory genes evaluated, with IL6, IL8, and PTGS2 varied the most between passage schemes (**Figure 6D**). Here, 91 out of 98 (93%) immunomodulatory fold-change comparisons differ by less than one order of magnitude (**Supplemental Table 6**), and 6 out of 14 (43%) immunomodulatory genes have low variability between passage schemes with 3 or fewer donors (less than 50%) showing significance differences in fold-change (**Supplemental Table 7**).

Finally, we observed that angiogenic gene expression was higher in male cells in the p4-5 passage scheme and higher in female cells in the p3-5 passage scheme (**Figure 6E**). Once again, we see that passage scheme impacts observed sex-based differences, and this is consistent with our passage scheme experiments where we saw that passage scheme significantly impacts cellular activity. Although only three genes were tested, only one angiogenic gene (VEGFA) displayed reversed sex-based differences based on passage scheme with female cells expressing significantly higher levels in the p3-5 scheme while male cells expressed significantly higher levels in the p4-5 scheme. The difference in gene expression fold changes between passage schemes was less than 2 for most of the angiogenic genes evaluated (**Figure 6F**). Here, 19 out of 21 (90%) angiogenic fold-change comparisons differ by less than one order of magnitude (**Supplemental Table 8**), and 1 out of 3 immunomodulatory genes have low variability between passage schemes with fewer than 3 donors (less than 50%) showing significant differences in fold-change (**Supplemental Table 9**). Notably, Donor G displayed the greatest differences in fold change between passage schemes in all genes with the greatest difference for VEGFA (**Figure 6E**). To summarize, these data show that male cells display overall enhanced osteogenic and immunomodulatory gene expression in the p4-5 scheme compared to female cells, with dramatic differences between passage scheme observed in donors E and G.

### 3.7. Male and female donors release similar levels of osteogenic factors, while male cells release significantly more immunomodulatory factors

We quantified osteogenic, immunomodulatory, extracellular matrix (ECM), and angiogenic factors secreted by all 8 donors throughout the 56-day culture period for p3-5 only using a custom Luminex panel (**Supplemental Figure 4-5**). For osteogenic factors, BMP2 was secreted at the highest levels by Day 56 followed by SPARC, OPN, and MMP13 (**Figure 7A**). No significant difference was observed between donors for osteogenic factors measured at Day 56. We observed that female cells secreted significantly more SPARC compared to male cells at Day 56; however, no other osteogenic factors displayed significant sex-differences (**Figure 7A**). For immunomodulatory factors, MMP2 was secreted at the highest levels by Day 56 followed by MMP3, IL6, GAL9, IL1B, and IL10 (**Figure 7B**). No significant difference was observed between donors for immunomodulatory factors measured at Day 56. Male cells displayed significantly higher immunomodulatory factor secretion compared to female cells, namely GAL9, IL10, IL1b, and MMP3 (**Figure 7B**). For extracellular matrix (ECM) factors, FN was expressed at the highest levels by Day 56 followed by IGFB3 (**Figure 8A**), and for angiogenic factors, VEGF was expressed at the highest levels by Day 56 compared to ANG (**Figure 8B**). No significant differences were observed between donors for ECM or angiogenic factors measured at Day 56. Male cells displayed significantly higher IGFB3 (ECM) and ANG (angiogenic) secretion compared to female cells (**Figure 8**). Overall, this cohort of male cells secreted significantly more immunomodulatory factors compared to this cohort of female cells, but there were no differences in osteogenic factor secretion between these male and female cells. To summarize, male donors display an enhance osteogenic and immunomodulatory soluble factor secretion compared to female cells at day 56.

**Figure 7:**
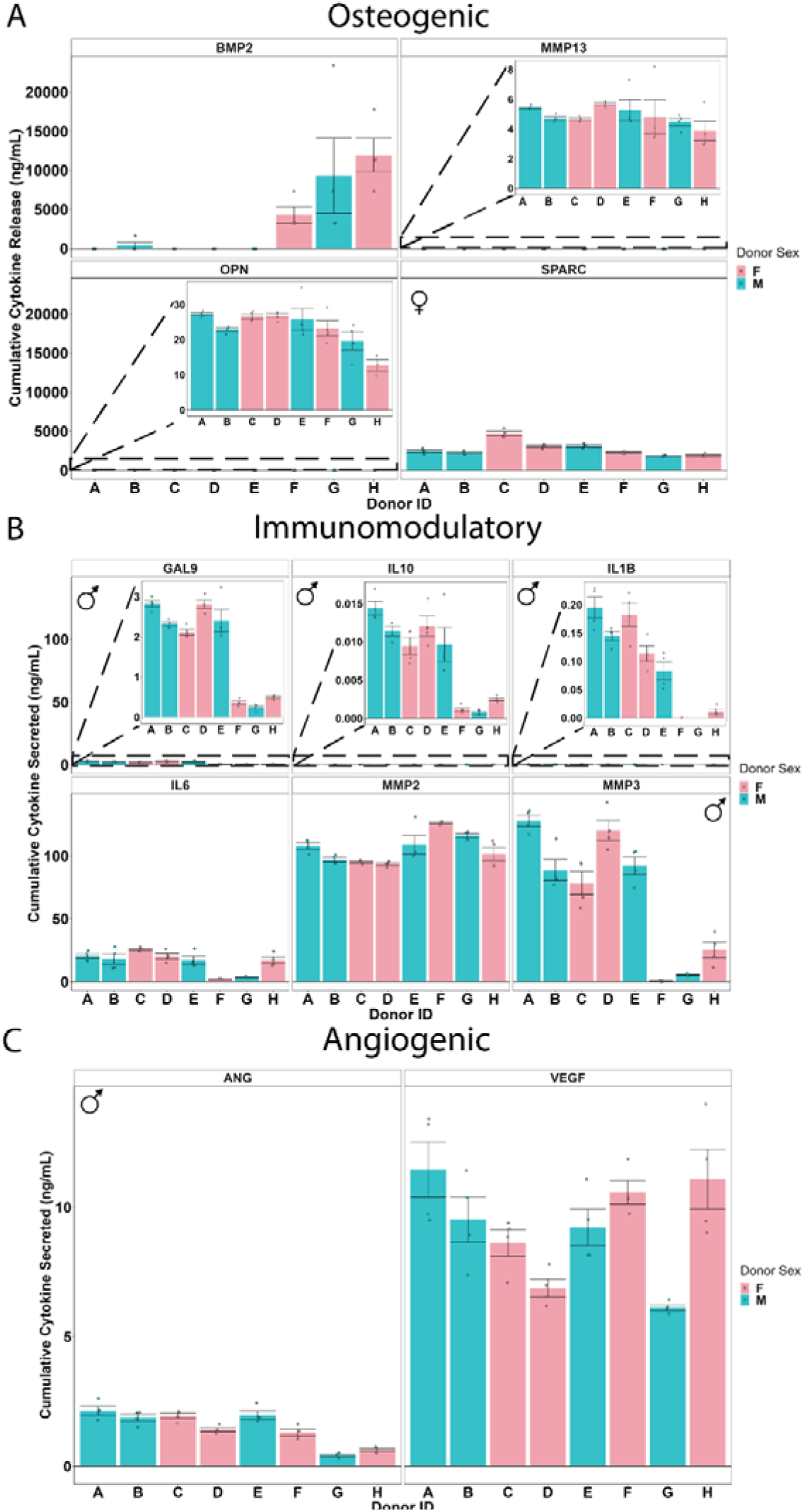
Osteogenic, immunomodulatory, and angiogenic protein secretion at Day 56. Eight donors (A, B, E, G – male, and C, D, F, H – female) with 2 passage methods (p4-5 and p3-5) were cultured on mineralized collagen scaffolds for 56-days. A Luminex assay was used to quantify secreted factors. (A) Female cells secrete more SPARC than male cells and there are no sex differences in the other osteogenic factors (B) Male cells secrete more immunomodulatory factors than female cells. (C) Male cells secrete significantly more ANG than female cells. Sex symbol (male: ♂, female: ♀) indicates sex-based significance (p<0.05) at the indicated timepoint. Cumulative cytokines secreted is plotted as mean +/- standard error.

## 4. Discussion

For the first time in our lab and in the field, we probed the influence of passage scheme on human mesenchymal stem cell (hMSC) response in 3D mineralized collagen scaffolds with well-defined osteogenic properties. Mesenchymal stem cells (MSCs) are used extensively in research across multiple disciplines for the development of novel therapeutics, tissue engineering solutions, and the understanding of drug efficacy [63]. Extensive work has been done to study how culture length and freeze-thaw cycles influence MSC metabolic activity on 2D substrates; however, little research exists about the impact of MSC passage scheme on key indicators of MSC viability and activity in 3D. We tested the influence of time between cell thawing and seeding and the final passage number on hMSC cell response over a 21-day culture period (**Figure 1A**). Beyond the influence of passage scheme, a more significant parameter that is largely absent and not reported in literature is the sex of donor cells [23]. Thus, we probed donor variability and sex-based differences on hMSC response (**Figure 1B**). Overall, it appears that despite differences in metabolic activity, cell number, and growth factor secretion (OPG) between passage schemes, functional outputs like alkaline phosphatase (ALP) activity, mineral deposition, and gene expression are comparable between passage schemes. Sex differences were observed at a higher frequency in passage scheme p4-5, indicating the importance of passage length on cellular outputs, but most of the significant sex-differences were consistent between passage schemes.

First, we compared the effect of passage scheme on commonly used metrics in the field for cell response and osteogenic potential (**Figure 2**). We compared three different culture groups on mineralized collagen scaffolds: cells thawed at passage 3 and seeded at passage 6 (p3-6), cells thawed at passage 3 and seeded at passage 5 (p3-5), and cells thawed at passage 4 and seeded at passage 5 (p4-5). Overall, we observed that metabolic activity and cell proliferation, commonly thought to be closely related, did not correlate across all donors. Therefore, cell metabolic activity and proliferation cannot be used interchangeably and should be reported for each donor should multiple donors be included in a study. Furthermore, we observed that p3-6 groups displayed the lowest metabolic activity for some donors (G – male, and F – female) and the lowest cell number for others (C – male, and F – female) and for one donor (G – male) did not produce sufficient cells for the full study (**Figure 2A,B**). This could be due to the final passage number at the time of seeding (passage 6 vs. passage 5) or the time cultured on 2D polystyrene before seeding, both of which have been shown to impact hMSC activity *in vitro* and *in vivo* [44, 64]. We further, measured osteogenic soluble factor secretion and observed significant differences in expression as a function of passage scheme (**Figure 2D**). Interestingly, we observed that donors that displayed the highest metabolic activity displayed some of the lowest soluble factor expression (B – male) further contrasting the commonly held notion of correlation between these metrics. These data suggest that passage scheme has a significant impact on cell outcomes and should be considered carefully when developing tissue engineering solutions. Further these data highlight a critical need to report detailed culture methods including cell passage at thawing and cell passage at experimental use.

Next, we evaluated donor variability and sex differences of cells seeded in our mineralized collagen scaffolds by quantifying their activity and their osteogenic, immunomodulatory, and angiogenic potential. We acquired cells from 8 donors (4 male, 4 female) to conduct this donor variability study using both adipose and bone marrow tissue (**Table 1**). We needed to use multiple companies to acquire this large donor set, due both to limited availability per company and limited availability of female donors for many companies. This illuminates a critical need for expanding the availability of cells from a variety of donors. Nonetheless, we compared these 8 donors cultured in p4-5 (thawed at passage 4 and seeded at passage 5) and p3-5 (thawed at passage 3 and seeded at passage 5) passage schemes and seeded them on mineralized collagen scaffolds for 56 days and quantified cell metabolic activity, proliferation, growth factor secretion, ALP activity, and gene expression.

We first evaluated donor and sex variability in cell metabolic activity and proliferation (**Figure 3**). In both passage schemes we observed male donors displaying significantly higher metabolic activity and cell proliferation compared to female donors. These differences occurred most often in passage scheme p4-5 further indicating that passage scheme has an impact on cell metabolic activity and cell number, and this is consistent with our passage scheme experiments (**Figure 2A**). Previous transcriptomic analysis of ASCs have observed sex-based differences suggesting potential influences on proliferation and differentiation with female cells having a lower potential than male cells [44] however others showed limited significance of sex on cell proliferation [65]. Taken together, we can conclude that this cohort of male cells is more metabolically active and proliferative than female cells on our mineralized collagen scaffolds. This would be an important consideration when using our scaffolds in patients, as the primary challenge in female patients would appear to be cell recruitment and proliferation; however, these might not be a challenge in male patients.

We then examined donor and sex variability in osteogenic potential via a combination of functional markers (OPG secretion; ALP activity; mineral deposition) and gene expression analysis (see **Supplemental Table 10-11** for order of magnitude differences and significance between donor variability assays). OPG is a decsuppoy receptor in the RANKL/RANK pathway inhibiting osteoclast formation and thus limiting bone resorption [66]. ALP is highly expressed in cells of mineralized tissue, facilitates mineralization, and is an indicator of bone metabolism [67]. We observed differences in OPG and ALP secretion between passage schemes suggesting the impact of passage scheme on downstream osteoclast activity and bone resorption (**Figure 4**) [66, 68]. The p4-5 groups displayed sex differences in ALP activity at later time points, with female cells having significantly higher ALP activity than male cells (**Figure 4C,D**). The variability observed in cell metabolic activity and cell number is not reflected ALP activity in this cohort of cells. We further evaluated mineral deposition at Day 56 and observed female donors deposit higher amounts of mineral than male cells in the p4-5 group (**Figure 5**). Interestingly the donor that displayed the lowest metabolic activity and cell number lead to the highest mineral deposition (Donor H – female). No sex differences were observed in mineral content in the p3-5 groups, but the mineral content fold change between p4-5 and p3-5 groups were similar. This suggests that cells are influencing mineral content similarly between passage schemes despite differences in metabolic activity and cell number. Next, we quantified osteogenic gene expression via a custom NanoString code set, where we observed male cells displaying significantly higher expression levels compared to female cells (**Figure 6** **and Supplemental Figures 1-3**). Most donors had similar osteogenic gene expression levels between passage schemes (difference in fold-change between passage schemes less than 2) with Donors E (male) and G (male) having the most variability between passage schemes. This aligns with previous findings form our lab that the mineralized collagen scaffolds provide significant instructive cues to promote osteogenesis *in vitro* [45, 48]. Finally, we quantified secreted osteogenic factor release via a Luminex assay, and observed female cells secreted significantly more SPARC than male cells by Day 56, but no sex differences were observed in the other osteogenic factors (**Figure 7A**). This suggests that this cohort of male and female cells have the same secretion levels of osteogenic factors. We have observed consistent osteogenic output on our scaffolds across species both *in vitro* [44, 45, 59, 69] and *in vivo* [50, 52, 54], and these results further confirm the efficacy of our mineralized collagen scaffolds for bone tissue engineering. We plan to work with our existing collaborators to conduct *in vivo* experiments to evaluate sex-differences in bone healing.

Additionally, we examined hMSC immunomodulatory and angiogenic potential via gene expression and soluble factors released. Male cells had significantly higher immunomodulatory gene expression compared to female cells in both passage schemes at Day 56 (**Figure 6C**). Immunomodulatory gene expression was the most variable group evaluated (**Figure 6D**), with several differences in fold-change greater than 2. Furthermore, male cells secreted significantly higher amounts of immunomodulatory cytokines (4 out of 6 analyzed cytokines: GAL9, IL10, IL1B, and MMP3) compared to female cells (**Figure 7B**) consistent with our gene expression data. This suggests there are significant sex-based differences in immunomodulatory potential of these hMSCs, and this has been observed *in vitro* [70, 71] by other groups. As seen in Table 1 various donors had co-morbidities while for others such health information was not provided. Thus, these differences in immunomodulatory potential could be attributed to different treatments, stress levels, and lifestyle habits that these patients experienced. In our previous work, we observed that stimulating hMSCs with inflammatory cytokines significantly enhances their immunomodulatory potential compared to a basal control [72] therefore, we would expect variability in hMSC immunomodulatory potential as a function of the systemic inflammatory state in patients. VEGFA gene expression was higher in male cells in the p4-5 passage scheme and higher in female cells in the p3-5 passage scheme with minimal variability in expression between passage schemes (**Figure 6E,F**). Male cells secreted significantly more VEGF compared to female cells by Day 56 (**Figure 7**). Given the limited number of angiogenic genes and growth factors characterized here, it is difficult to make meaningful conclusions; therefore, further analysis would need to be done to characterize the impact of donor and sex on angiogenic potential.

Overall, we observe that passage number has little influence on osteogenic potential (ALP activity, mineral content) but rather we observed critical sex-based differences. Altogether, these data suggest that female cells possess a higher (p4-5) or equivalent (p3-5) osteogenic capacity in mineralized collagen scaffolds compared to male cells despite displaying lower metabolic activity and proliferation than male cells (p4-5 and p3-5). Thus, there may be a difference between male and female patient needs when it comes to therapeutic intervention, where male patients might require a greater boost in osteogenic activity while female patients might require an enhancement in their cell proliferation and migration capacity. A limitation of this study, and an important consideration when studying sex differences, is the consideration of sex hormones. Estrogen is known to play vital roles in inhibiting bone resorption [73] but has also been shown to interact with the immune system [74]. Although in this work we did not directly study the role of estrogens and other sex hormones, we conducted these experiments in phenol-red free media because phenol-red is known to be a weak estrogen that can stimulate estrogen sensitive cells [75]. However, future studies should expand these efforts and study sex-based differences in hormone-stripped media as serum additives provide significant hormone levels to the system.

These 56-day experiments are time consuming, large, and costly, and not all research labs have the resources needed to conduct such large experiments. The scale and cost of these experiments is a primary deterrent in incorporating multiple donors in an experiment and considering sex as an experimental variable. Ongoing efforts in our lab are focused on developing high-throughput platforms to allow for the rapid evaluation of donor variability and donor characteristics as biological variables in 3D mineralized collagen biomaterials. Further, we are interested in how tissue engineers can repurpose current high throughput technologies to evaluate donor variability *in vitro*. Finally, sourcing enough cells from a large pool of donors is an additional challenge in conducting these experiments. Thus, this work is a call to action for academics and industry to expand their donor pools across sex, age, race, and pre-existing conditions and make these cells more accessible to researchers. This access to diverse pools of donor cells would allow for the broadening of knowledge and reporting of findings considering subpopulations of people without generalizations that could lead to more harm.

## 5. Conclusions

Human mesenchymal stem cell (hMSC) passage number and culture time post thawing are important variables to determine hMSC osteogenic response, and it is essential that these are reported accurately. Furthermore, donor variability and donor sex are important factors when designing biomaterials for bone repair to ensure the results are applicable to a wide range of potential patients. We want to ensure our mineralized collagen scaffolds can heal a diverse set of patients with varying age, sex, race/ethnicity, and health status. First, we evaluated differences in hMSC response as a function of passage number and culture time post thawing. Our findings show that final passage number does not elicit a consistent response regardless of culture time with significantly different functional responses observed; however, in our donor variability experiments genomic trends appear to be consistent between passage schemes. Therefore, it is essential that these culture parameters are reported in literature to allow for comparisons of data across labs. Then, we evaluated donor variability between 8 donors cultured on mineralized collagen scaffolds. An osteogenic response was observed for all donors on our mineralized collagen scaffolds, showing their applicability for a large pool of potential patients. We report significantly increased metabolic activity and proliferation for male cells, and significantly higher alkaline phosphate (ALP) activity, osteoprotegerin (OPG) secretion, and mineral deposition in female donors. Mineralized collagen scaffolds provide a platform to study donor variability and use donor characteristics such as sex as experimental variables. Our work illuminates the need to expand the literature by studying donor variability and reporting patient characteristics to allow for transparency of data and results.

## Acknowledgements

The authors would like to acknowledge the following institutes for access to their facilities and services: the School of Chemical Sciences Microanalysis Laboratory, the Carl R. Woese Institute for Genomic Biology, the Tumor Engineering and Phenotyping Shared Resource (TEP) at the Cancer Center at Illinois, and the Beckman Institute for Advanced Science and Technology, located at the University of Illinois. Research reported in this publication was supported by the National Institute of Dental and Craniofacial Research of the National Institutes of Health under Award Number R21 DE026582 and R01 DE030491 (BACH) as well as National Institute of Arthritis and Musculoskeletal and Skin Diseases under Award Number R01 AR077858 (BACH). We are also grateful for funds provided by the NSF Graduate Research Fellowship (DGE-1746047 to VK; DGE-1144245 to AST) and the Chemistry-Biology Interface Research Training Program at the University of Illinois (T32 GM070421, VK). Additional support was provided by the Carl R. Woese Institute for Genomic Biology and the Chemical and Biomolecular Engineering Dept. at the University of Illinois at Urbana-Champaign. The interpretations and conclusions presented are those of the authors and are not necessarily endorsed by the National Institutes of Health or the National Science Foundation.

## Contributions (CRediT: Contributor Roles Taxonomy [51, 52])

**Vasiliki Kolliopoulos:** Conceptualization, Data curation, Formal Analysis, Visualization, Investigation, Methodology, Writing – original draft, Writing – review & editing. **Aleczandria Tiffany:** Conceptualization, Data curation, Formal Analysis, Visualization, Investigation, Methodology, Writing – original draft, Writing – review & editing. **Maxwell Polanek:** Investigation, Data curation, Formal Analysis. **Brendan Harley:** Conceptualization, Resources, Project administration, Funding acquisition, Supervision, Writing – review & editing.

## Data availability

The raw data required to reproduce these findings are available upon request to Brendan Harley. The processed data required to reproduce these findings are available upon request to Brendan Harley.

