## Supplemental Data for "DONOR VARIABILITY IN HUMAN MESENCHYMAL STEM CELL OSTEOGENIC RESPONSE AS A FUNCTION OF PASSAGE CONDITIONS AND DONOR SEX"

[1] Dept. Chemical and Biomolecular Engineering, University of Illinois at Urbana-Champaign

Urbana, IL 61801

[2] Cancer Center at Illinois, University of Illinois at Urbana-Champaign

Urbana, IL 61801

[3] Carl R. Woese Institute for Genomic Biology, University of Illinois at Urbana-Champaign

Urbana, IL 61801

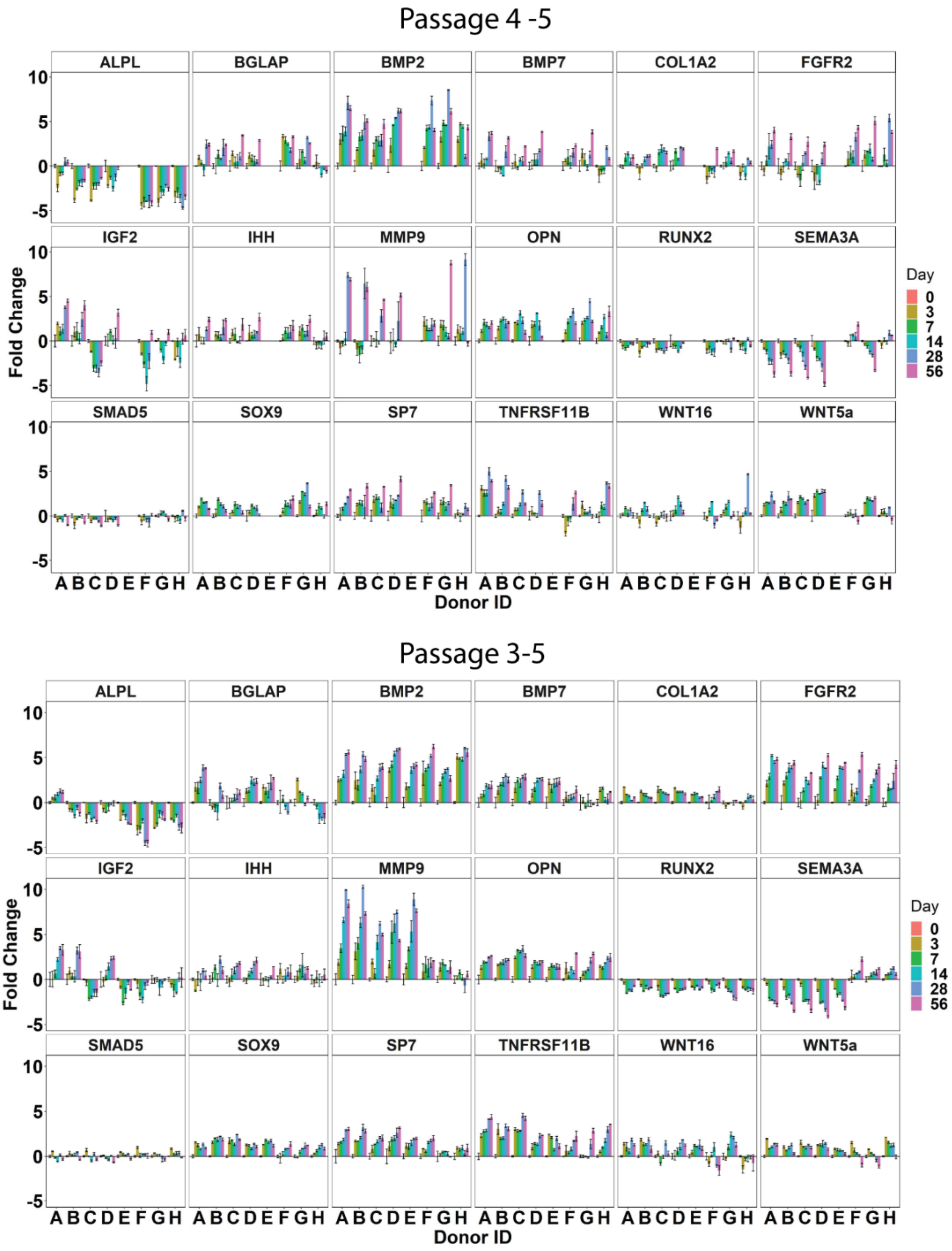


**Supplemental Figure 1: Osteogenic gene expression of 8 donors cultured on mineralized collagen scaffolds.** Eight donors (A, B, E, G – male, and C, D, F, H – female) with 2 passage methods (p4-5 and p3-5) were cultured on mineralized collagen scaffolds for 56-days. Osteogenic gene expression was quantified using NanoString at days 3, 7, 14, 28, and 56 and normalized to a Day 0 control. Fold change is plotted as the mean +/- standard error.

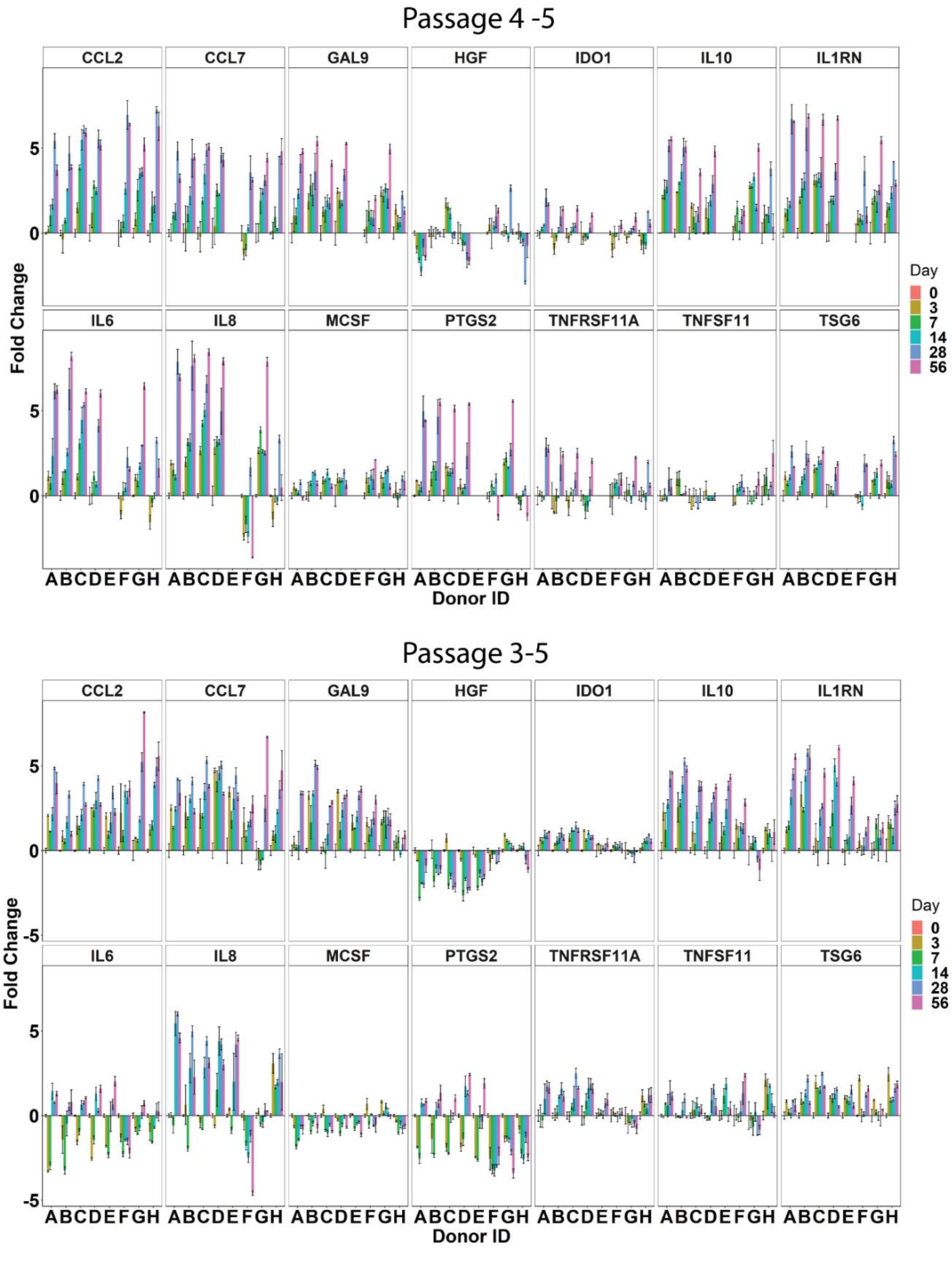


**Supplemental Figure 2: Immunomodulatory gene expression of 8 donors cultured on mineralized collagen scaffolds.** Eight donors (A, B, E, G – male, and C, D, F, H – female) with 2 passage methods (p4-5 and p3-5) were cultured on mineralized collagen scaffolds for 56-days. Immunomodulatory gene expression was quantified using NanoString at days 3, 7, 14, 28, and 56 and normalized to a Day 0 control. Fold change is plotted as the mean +/- standard error.

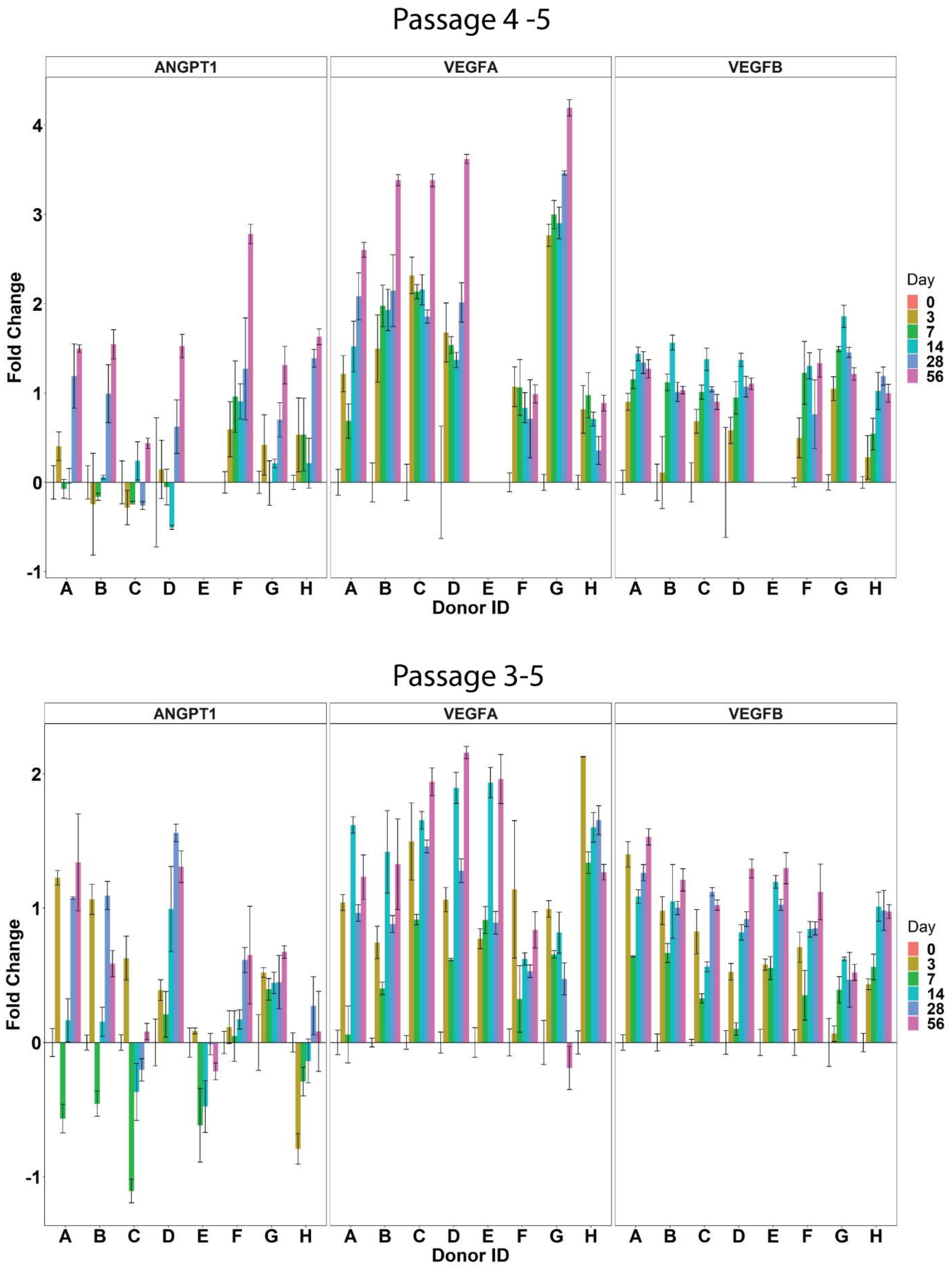


**Supplemental Figure 3: Angiogenic gene expression of 8 donors cultured on mineralized collagen scaffolds.** Eight donors (A, B, E, G – male, and C, D, F, H – female) with 2 passage methods (p4-5 and p3-5) were cultured on mineralized collagen scaffolds for 56-days. Angiogenic gene expression was quantified using NanoString at days 3, 7, 14, 28, and 56 and normalized to a Day 0 control. Fold change is plotted as the mean +/- standard error.

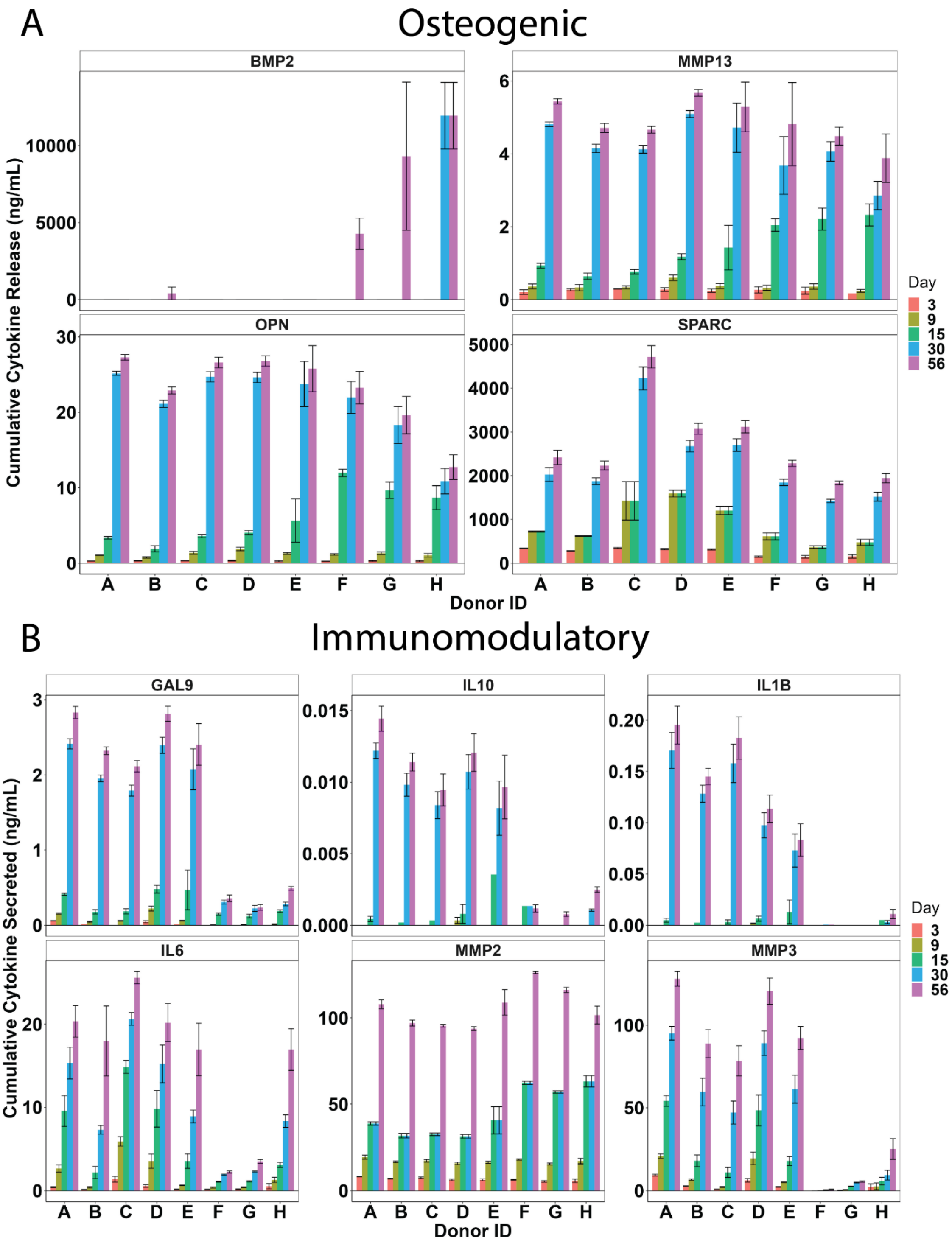


**Supplemental Figure 4: Osteogenic and Immunomodulatory secreted factor release of 8 donors on mineralized collagen scaffolds.** Eight donors (A, B, E, G – male, and C, D, F, H – female) with 2 passage methods (p4-5 and p3-5) were cultured on mineralized collagen scaffolds for 56-days. Osteogenic and immunomodulatory secreted factors were quantified using a Luminex assay. Media was pooled and analyzed for each replicate (n=6) at days 3, 9, 15, 30, and 56. Cumulative cytokine secretion is plotted as the mean +/- standard error.

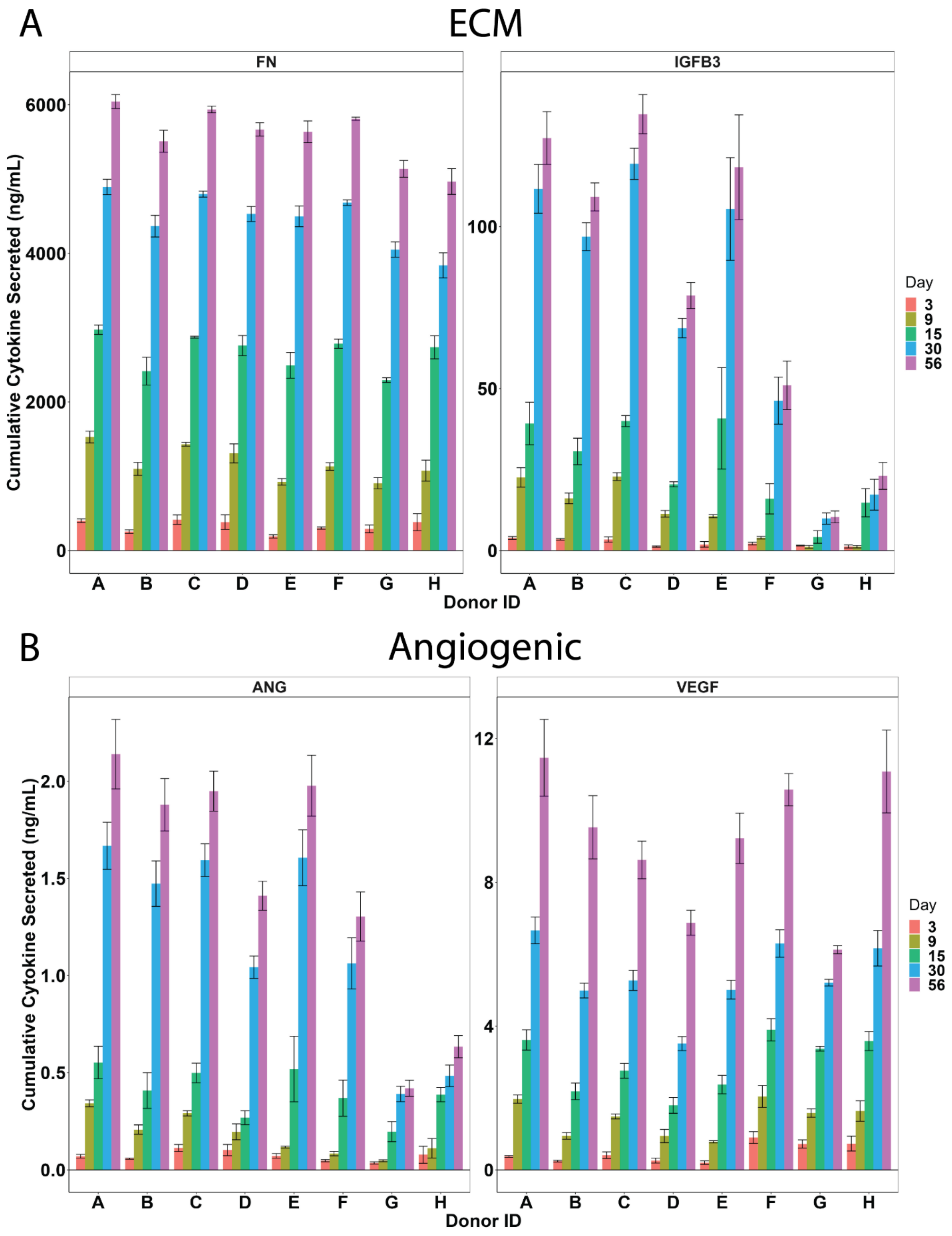


**Supplemental Figure 5: Matrix and angiogeneic secreted factor release of 8 donors on mineralized collagen scaffolds.** Eight donors (A, B, E, G – male, and C, D, F, H – female) with 2 passage methods (p4-5 and p3-5) were cultured on mineralized collagen scaffolds for 56-days. Extracellular matrix (ECM) and angiogenic secreted factors were quantified using a Luminex assay. Media was pooled and analyzed for each replicate (n=6) at days 3, 9, 15, 30, and 56. Cumulative cytokine secretion is plotted as the mean +/- standard error.

**Supplemental Table 1: NanoString gene panel.**

| Customer Name | HUGO Gene | Full Name | Probe NSID | Category |
|---|---|---|---|---|
| ALPL | ALPL | Alkaline Phosphatase | NM_000478.3:2065 | Osteogenic |
| ANGPT1 | ANGPT1 | Angiopoietin 1 | NM_001146.3:2080 | Angiogenic |
| BGLAP | BGLAP | Bone Gamma-Carboxyglutamate Protein | NM_199173.4:44 | Osteogenic |
| BMP2 | BMP2 | Bone Morphogenic Protein 2 | NM_001200.2:1515 | Osteogenic |
| BMP7 | BMP7 | Bone Morphogenic Protein 7 | NM_001719.1:525 | Osteogenic |
| CCL2 | CCL2 | C-C Motif Chemokine Ligand 2 | NM_002982.3:123 | Immunomodulatory |
| CCL7 | CCL7 | C-C Motif Chemokine Ligand 7 | NM_006273.2:120 | Immunomodulatory |
| COL1A2 | COL1A2 | Collagen Type I Alpha 2 Chain | NM_000089.3:2635 | Osteogenic |
| MCSF | CSF1 | Colony Stimulating Factor 1 | NM_000757.4:823 | Immunomodulatory |
| IL8 | CXCL8 | Interleukin 8 | NM_000584.2:25 | Immunomodulatory |
| FGFR2 | FGFR2 | Fibroblast Growth Factor Receptor 2 | NM_000141.4:2204 | Osteogenic |
| GAPDH | GAPDH | Glyceraldehyde-3-Phosphate Dehydrogenase | NM_001256799.1:386 | Housekeeping |
| GUSB | GUSB | Glucuronidase Beta | NM_000181.3:1899 | Housekeeping |
| HGF | HGF | Hepatocyte Growth Factor | NM_000601.4:550 | Immunomodulatory |
| IDO1 | IDO1 | Indoleamine 2,3-Dioxygenase 1 | NM_002164.5:369 | Immunomodulatory |
| IGF2 | IGF2 | Insulin Like Growth Factor 2 | NM_000612.4:765 | Osteogenic |
| IHH | IHH | Indian Hedgehog Signaling Molecule | NM_002181.2:1693 | Osteogenic |
| IL10 | IL10 | Interleukin 10 | NM_000572.2:622 | Immunomodulatory |
| IL1RN | IL1RN | Interleukin 1 Receptor Antagonist | NM_000577.3:480 | Immunomodulatory |
| IL6 | IL6 | Interleukin 6 | NM_000600.3:364 | Immunomodulatory |
| GAL9 | LGALS9 | Galectin 9 | NM_002308.3:359 | Immunomodulatory |
| MMP9 | MMP9 | Matrix Metallopeptidase 9 | NM_004994.2:1530 | Osteogenic |
| OAZ1 | OAZ1 | Ornithine Decarboxylase Antizyme 1 | NM_004152.2:313 | Housekeeping |
| PTGS2 | PTGS2 | Prostaglandin-Endoperoxide Synthase 2 | NM_000963.1:495 | Immunomodulatory |
| RUNX2 | RUNX2 | RUNX Family Transcription Factor 2 | NM_004348.3:1850 | Osteogenic |
| SEMA3A | SEMA3A | Semaphorin 3A | NM_006080.1:585 | Osteogenic |
| SMAD5 | SMAD5 | SMAD Family Member 5 | NM_005903.5:1044 | Osteogenic |
| SOX9 | SOX9 | SRY-Box Transcription Factor 9 | NM_000346.2:2135 | Osteogenic |
| SP7 | SP7 | Sp7 Transcription Factor (Osterix) | NM_001173467.1:1510 | Osteogenic |
| OPN | SPP1 | Secreted Phosphoprotein 1 (Osteopontin) | NM_000582.2:760 | Osteogenic |
| TSG6 | TNFAIP6 | TNF-Stimulated Gene 6 Protein | NM_007115.2:250 | Immunomodulatory |
| TNFRSF11A | TNFRSF11A | TNF Receptor Superfamily Member 11a (RANK) | NM_003839.3:226 | Immunomodulatory |
| TNFRSF11B | TNFRSF11B | Osteoprotegerin | NM_002546.2:1075 | Osteogenic |
| TNFSF11 | TNFSF11 | TNF Superfamily Member 11 (RANKL) | NM_003701.2:490 | Immunomodulatory |
| VEGFA | VEGFA | Vascular Endothelial Growth Factor A | NM_001025366.1:1325 | Angiogenic |
| VEGFB | VEGFB | Vascular Endothelial Growth Factor B | NM_003377.3:687 | Angiogenic |
| WNT16 | WNT16 | Wnt Family Member 16 | NM_057168.1:1621 | Osteogenic |
| WNT5a | WNT5A | Wnt Family Member 5A | NM_003392.3:475 | Osteogenic |

**Supplemental Table 2: Luminex soluble factor list.**

| Category | Soluble factor |
|---|---|
| Osteogenic | BMP2 |
| | SPARC |
| | Osteopontin (OPN) |
| | IGFb3 |
| | Fibronectin |
| | MMP13 |
| Immunomodulatory | IL1b |
| | IL6 |
| | Galectin9 |
| | IL10 |
| | MMP2 |
| | MMP3 |
| Angiogenic | Angiogenin |
| | VEGF |

**Supplemental Table 3: Sex differences in gene expression at Day 56 for donor variability experiments.** We evaluated sex differences in osteogenic, immunomodulatory, and angiogenic gene expression in passage schemes p4-5 and p3-5 at Day 56. A value p<0.05 indicates significantly different expression between male and female cells. 'Male' and 'Female' indicate the sex of donor cells that had significantly higher expression. NS indicates no significance in expression between male and female cells.

Sex differences occurred more frequently in p4-5 (25 out of 35 total genes) compared to p3-5 (16 out of 35 total genes). Male cells had significantly higher expression of osteogenic and immunomodulatory genes in both p4-5 (12 out of 18 osteogenic genes, 8 out of 14 immunomodulatory genes) and p3-5 (5 out of 18 osteogenic genes, 5 out of 14 immunomodulatory genes).

| | | **p4-5 p-value** | **p4-5 sex significance** | **p3-5 p-value** | **p3-5 sex significance** |
|---|---|---|---|---|---|
| **Osteogenic** | ALPL | 6.95E-06 | Male | 6.04E-06 | Male |
| | BGLAP | 0.034 | Male | 1.77E-06 | Male |
| | BMP2 | 2.20E-04 | Male | 5.15E-05 | Female |
| | BMP7 | 1.82E-08 | Male | 0.103 | NS |
| | COL1A2 | 0.179 | NS | 0.003 | Female |
| | FGFR2 | 0.007 | Male | 0.510 | NS |
| | IGF2 | 5.18E-07 | Male | 0.010 | Male |
| | IHH | 0.045 | Male | 0.370 | NS |
| | MMP9 | 8.94E-12 | Male | 2.32E-10 | Male |
| | OPN | 0.048 | Male | 0.160 | NS |
| | RUNX2 | 0.001 | Male | 0.704 | NS |
| | SEMA3A | 2.15E-09 | Female | 1.07E-06 | Female |
| | SMAD5 | 0.137 | NS | 0.333 | NS |
| | SOX9 | 0.254 | NS | 0.934 | NS |
| | SP7 | 0.001 | Male | 0.655 | NS |
| | TNFRSF11B | 0.259 | NS | 0.266 | NS |
| | WNT16 | 0.648 | NS | 0.001 | Male |
| | WNT5a | 1.49E-06 | Male | 0.874 | NS |
| **Immunomodulatory** | CCL2 | 6.79E-05 | Female | 0.529 | NS |
| | CCL7 | 0.322 | NS | 0.511 | NS |
| | GAL9 | 6.18E-09 | Male | 1.31E-05 | Male |
| | HGF | 0.452 | NS | 0.003 | Male |
| | IDO1 | 0.005 | Male | 0.351 | NS |
| | IL8 | 1.18E-10 | Male | 0.001 | Male |
| | IL10 | 1.66E-07 | Male | 0.300 | NS |
| | IL1RN | 6.81E-08 | Male | 0.249 | NS |
| | IL6 | 3.75E-10 | Male | 0.001 | Male |
| | MCSF | 0.002 | Female | 0.915 | NS |
| | PTGS2 | 1.33E-13 | Male | 0.023 | Male |
| | TNFRSF11A | 4.41E-05 | Male | 0.056 | NS |
| | TSG6 | 0.027 | Female | 4.35E-06 | Female |
| | TNFSF11 | 0.766 | NS | 0.002 | Female |
| **Angiogenic** | ANGPT1 | 0.165 | NS | 0.675 | NS |
| | VEGFA | 1.76E-11 | Male | 0.001 | Female |
| | VEGFB | 0.238 | NS | 0.610 | NS |

**Supplemental Table 4: Order of magnitude difference in osteogenic gene expression between donor variability experiments at Day 56.** A value equal to 1 indicates a one order of magnitude (OOM) difference in expression between passage schemes p4-5 and p3-5. All values greater than 1 are red and italicized, and all values that round to 1 (i.e., greater than 0.5) are bolded and italicized. Donors A, B, and G are male. Donors C, D, F, and H are female. Donor E was not analyzed in p4-5 due to lack of cells available at cell seeding and therefore not present in this table.

117 out of 126 (93%) osteogenic fold change comparisons differ by less than one OOM between passage schemes. Donor G (male) is the most variable between passage schemes: 4 out of 18 osteogenic genes have OOM differences greater than 1 and 5 out of 18 osteogenic genes have OOM differences greater than 0.5. Donors C and H (both female) are the least variable between passage schemes: 1 out of 18 osteogenic genes has an OOM difference greater than 0.5.

| | order of magnitude difference in fold change for osteogenic genes | | | | | | |
|---|---|---|---|---|---|---|---|
| ***ALPL*** | 0.385 | 0.104 | 0.191 | 0.322 | 0.036 | 0.136 | 0.092 |
| ***BGLAP*** | 0.200 | 0.435 | 0.491 | 0.077 | *1.313* | ***0.676*** | 0.444 |
| ***BMP2*** | 0.064 | 0.022 | 0.072 | 0.016 | 0.185 | 0.360 | 0.109 |
| ***BMP7*** | 0.273 | 0.103 | 0.112 | 0.167 | 0.206 | *1.283* | 0.142 |
| ***COL1A2*** | 0.262 | 0.382 | 0.237 | 0.306 | 0.129 | ***0.880*** | 0.071 |
| ***FGFR2*** | 0.080 | 0.130 | 0.081 | 0.338 | 0.093 | 0.103 | 0.043 |
| ***IGF2*** | 0.140 | 0.109 | 0.227 | 0.122 | 0.380 | *1.301* | 0.477 |
| ***IHH*** | ***0.732*** | 0.359 | 0.007 | 0.094 | 0.335 | 0.287 | 0.042 |
| ***MMP9*** | 0.086 | 0.082 | 0.032 | 0.074 | 0.022 | ***0.804*** | 0.254 |
| ***OPN*** | 0.104 | 0.024 | 0.443 | ***0.638*** | 0.158 | 0.120 | 0.129 |
| ***RUNX2*** | 0.338 | ***0.501*** | 0.244 | ***0.620*** | *1.525* | ***0.909*** | 0.370 |
| ***SEMA3A*** | 0.121 | 0.020 | 0.077 | 0.067 | 0.069 | 0.443 | 0.014 |
| ***SMAD5*** | 0.433 | 0.301 | 0.458 | 0.197 | 0.263 | 0.061 | 0.352 |
| ***SOX9*** | 0.066 | ***0.501*** | ***0.690*** | *1.029* | 0.158 | 0.065 | 0.202 |
| ***SP7*** | 0.019 | 0.079 | 0.208 | 0.118 | 0.110 | *1.693* | 0.099 |
| ***TNFRSF11B*** | 0.045 | 0.021 | 0.496 | 0.226 | 0.073 | *1.757* | 0.025 |
| ***WNT16*** | *1.259* | *1.426* | 0.125 | 0.453 | 0.491 | ***0.788*** | 0.407 |
| ***WNT5a*** | 0.089 | ***0.797*** | 0.254 | ***0.532*** | 0.129 | 0.249 | ***0.979*** |
| | ***A*** | ***B*** | ***C*** | ***D*** | ***F*** | ***G*** | ***H*** |

**Supplemental Table 5: Significance in osteogenic gene expression between donor variability experiments at Day 56.** A value $p<0.05$ indicates significantly different expression between passage schemes p4-5 and p3-5. All $p<0.05$ are red and italicized in the table. Donors A, B, and G are male. Donors C, D, F, and H are female. Donor E was not analyzed in p4-5 due to lack of cells available at cell seeding and therefore not present in this table.

12 out of 18 (67%) osteogenic genes have low variability between passage schemes with 3 or fewer donors (less than 50%) showing significant differences in fold-change. These 12 gene names are bolded and italicized in the table: ALPL, BMP2, BMP7, FGFR2, IGF2, IHH, MMP9, SEMA3A, SOX9, SP7, TNFRSF11B, and WNT16. SP7 has the least variable expression between passage schemes: 1 donor (G, male) has significantly different expression.

Donor G (male) is the most variable between passage schemes: 13 out of 18 osteogenic genes have significantly different expression. Donor H (female) is the least variable between passage schemes: 0 out of 18 osteogenic genes have significantly different expression.

| | **p-values for osteogenic gene expression between p4-5 and p3-5** | | | | | | |
|---|---|---|---|---|---|---|---|
| ***ALPL*** | *0.038* | 0.199 | *0.030* | 0.776 | 0.481 | 0.066 | 0.351 |
| BGLAP | *0.005* | 0.056 | *0.008* | 0.250 | *1.62E-04* | *0.002* | 0.168 |
| ***BMP2*** | 0.061 | 0.508 | 0.296 | 0.425 | *0.002* | *0.002* | 0.083 |
| ***BMP7*** | *0.034* | 0.172 | 0.178 | *0.013* | 0.126 | *0.003* | 0.088 |
| COL1A2 | 0.103 | *0.017* | *0.037* | *0.003* | 0.214 | *0.006* | 0.891 |
| ***FGFR2*** | 0.114 | 0.051 | 0.393 | *0.001* | *0.027* | 0.115 | 0.456 |
| ***IGF2*** | 0.165 | 0.385 | 0.119 | 0.166 | *0.011* | *0.044* | 0.762 |
| ***IHH*** | *0.037* | *0.047* | 0.971 | 0.396 | 0.393 | 0.076 | 0.950 |
| ***MMP9*** | *0.049* | 0.118 | 0.138 | *0.042* | 0.786 | *0.004* | 0.064 |
| OPN | 0.087 | 0.489 | *0.005* | *3.10E-04* | *0.010* | *0.022* | 0.311 |
| RUNX2 | 0.062 | *0.001* | *0.041* | *0.005* | 0.135 | *0.001* | 0.104 |
| ***SEMA3A*** | 0.070 | 0.561 | *0.043* | 0.070 | 0.360 | *4.65E-06* | 0.913 |
| SMAD5 | *0.005* | *0.039* | *0.005* | *0.008* | 0.892 | 0.763 | 0.460 |
| ***SOX9*** | 0.228 | *0.004* | *0.008* | *0.005* | 0.152 | 0.401 | 0.138 |
| ***SP7*** | 0.531 | 0.162 | 0.060 | 0.075 | 0.162 | *2.19E-04* | 0.704 |
| ***TNFRSF11B*** | 0.254 | 0.631 | *0.002* | 0.105 | 0.546 | *0.001* | 0.510 |
| ***WNT16*** | *0.0425* | 0.123 | 0.364 | *0.031* | 0.181 | *0.043* | 0.367 |
| WNT5a | 0.1782 | *0.001* | *0.024* | *0.001* | 0.434 | *1.18E-04* | 0.259 |
| | ***A*** | ***B*** | ***C*** | ***D*** | ***F*** | ***G*** | ***H*** |

**Supplemental Table 6: Order of magnitude difference in immunomodulatory gene expression between donor variability experiments at Day 56.** A value equal to 1 indicates a one order of magnitude (OOM) difference in expression between passage schemes p4-5 and p3-5. All values greater than 1 are red and italicized, and all values that round to 1 (i.e., greater than 0.5) are bolded and italicized. Donors A, B, and G are male. Donors C, D, F, and H are female. Donor E was not analyzed in p4-5 due to lack of cells available at cell seeding and therefore not present in this table.

91 out of 98 (93%) immunomodulatory fold-change comparisons differ by less than one OOM between passage schemes. Donor G (male) is the most variable between passage schemes: 2 out 14 immunomodulatory genes have OOM differences greater than 1 and 5 out of 14 immunomodulatory genes have OOM differences greater than 0.5. Donors A (male) and D (female) were the least variable between passage schemes: 0 out of 14 immunomodulatory genes have OOM differences greater than 1 and 3 out of 14 immunomodulatory genes have OOM differences greater than 0.5.

| | **order of magnitude difference in fold change for immunomodulatory genes** | | | | | | |
|---|---|---|---|---|---|---|---|
| ***CCL2*** | 0.032 | ***0.592*** | 0.336 | 0.280 | 0.242 | 0.195 | 0.052 |
| ***CCL7*** | 0.024 | 0.285 | 0.127 | 0.107 | 0.061 | 0.180 | 0.006 |
| ***GAL9*** | 0.153 | 0.042 | 0.156 | 0.194 | 0.168 | 0.434 | 0.155 |
| ***HGF*** | 0.208 | *1.273* | *1.283* | 0.138 | ***0.712*** | 0.328 | 0.184 |
| ***IDO1*** | 0.190 | 0.276 | 0.085 | 0.127 | ***0.554*** | *1.222* | 0.017 |
| ***IL8*** | 0.082 | 0.023 | 0.027 | 0.104 | 0.335 | ***0.641*** | 0.464 |
| ***IL10*** | 0.071 | 0.096 | 0.161 | 0.045 | 0.255 | ***0.612*** | 0.029 |
| ***IL1RN*** | ***0.680*** | *1.004* | ***0.764*** | ***0.576*** | 0.150 | ***0.946*** | ***0.830*** |
| ***IL6*** | 0.182 | ***0.550*** | 0.434 | 0.421 | 0.102 | *1.466* | ***0.615*** |
| ***MCSF*** | ***0.714*** | 0.006 | 0.229 | 0.019 | ***0.577*** | ***0.992*** | 0.138 |
| ***PTGS2*** | ***0.692*** | ***0.608*** | ***0.691*** | 0.347 | 0.278 | 0.213 | 0.303 |
| ***TNFRSF11A*** | 0.226 | 0.325 | 0.185 | 0.111 | ***0.628*** | 0.449 | 0.294 |
| ***TSG6*** | 0.343 | 0.336 | *1.750* | ***0.980*** | ***0.842*** | 0.076 | *1.080* |
| ***TNFSF11*** | 0.436 | 0.470 | 0.197 | ***0.552*** | 0.052 | ***0.755*** | 0.123 |
| | ***A*** | ***B*** | ***C*** | ***D*** | ***F*** | ***G*** | ***H*** |

**Supplemental Table 7: Significance in immunomodulatory gene expression between donor variability experiments at Day 56.** A value $p<0.05$ indicates significantly different expression between passage schemes p4-5 and p3-5. All $p<0.05$ are red and italicized in the table. Donors A, B, and G are male. Donors C, D, F, and H are female. Donor E was not analyzed in p4-5 due to lack of cells available at cell seeding and therefore not present in this table. 6/14 immunomodulatory genes have low variability between passage schemes with less than 50% of donors showing significant differences in fold-change. These 6 gene names are bolded and italicized in the table.

6 out of 14 (43%) immunomodulatory genes have low variability between passage schemes with 3 or fewer donors (less than 50%) showing significant differences in fold-change. These 6 gene names are bolded and italicized in the table: CCL7, HGF, IDO1, IL10, TNFRSF11A, and TNFSF11. IDO1, TNFRSF11A and TNFSF11 have the least variable expression between passage schemes: 2 out of 7 donors have significantly different expression.

Donor G (male) is the most variable between passage schemes: 12 out of 14 immunomodulatory genes have significantly different expression. Donor H (female) is the least variable between passage schemes: 1 out of 14 immunomodulatory genes has significantly different expression.

| | **p-values for immunomodulatory gene expression between p4-5 and p3-5** | | | | | | |
|---|---|---|---|---|---|---|---|
| CCL2 | 0.714 | *1.05E-04* | *0.002* | *0.009* | *0.023* | *0.015* | 0.583 |
| ***CCL7*** | 0.833 | *0.001* | *0.011* | 0.127 | 0.506 | *0.010* | 0.964 |
| GAL9 | *0.002* | 0.190 | *0.009* | *0.004* | *0.041* | *0.003* | 0.261 |
| ***HGF*** | 0.297 | *0.023* | *0.005* | 0.086 | *0.048* | 0.630 | 0.635 |
| ***IDO1*** | *0.010* | *0.029* | 0.285 | 0.152 | 0.362 | 0.056 | 0.929 |
| IL8 | *0.004* | *0.025* | *2.95E-04* | *3.71E-04* | *0.015* | *0.001* | 0.477 |
| ***IL10*** | *0.004* | 0.522 | 0.527 | 0.065 | *0.015* | *0.005* | 0.530 |
| IL1RN | *0.018* | 0.186 | *0.008* | *0.018* | 0.212 | *0.001* | 0.742 |
| IL6 | *2.22E-04* | *0.005* | *1.70E-04* | *1.28E-04* | *0.002* | *4.16E-05* | 0.141 |
| MCSF | 0.104 | *0.011* | *0.001* | *0.020* | *0.024* | *0.040* | 0.097 |
| PTGS2 | *3.88E-04* | *0.005* | *1.07E-04* | *7.79E-06* | 0.145 | *4.27E-04* | *0.014* |
| ***TNFRSF11A*** | 0.052 | *0.032* | 0.078 | 0.248 | 0.158 | *0.005* | 0.309 |
| TSG6 | 0.098 | *0.010* | *0.022* | *0.002* | 0.311 | *0.008* | 0.088 |
| ***TNFSF11*** | 0.305 | 0.281 | 0.472 | 0.305 | *2.38E-04* | *0.019* | 0.075 |
| | ***A*** | ***B*** | ***C*** | ***D*** | ***F*** | ***G*** | ***H*** |

**Supplemental Table 8: Order of magnitude difference in angiogenic gene expression between passage schemes at Day 56.** A value equal to 1 indicates a one order of magnitude (OOM) difference in expression between passage schemes p4-5 and p3-5. All values greater than 1 are red and italicized, and all values that round to 1 (i.e., greater than 0.5) are bolded and italicized. Donors A, B, and G are male. Donors C, D, F, and H are female. Donor E was not analyzed in p4-5 due to lack of cells available at cell seeding and therefore not present in this table.

19 out of 21 (90%) angiogenic fold-change comparisons differ by less than one OOM between passage schemes. Donors G (male) and H (female) are the most variable: 1 out of 3 angiogenic genes have OOM differences greater than 1 and 0 out of 3 angiogenic genes have OOM differences greater than 0.5. Donors A (male), B (male), and D (female) are the least variable: all OOM differences are less than 0.5.

| | **order of magnitude difference in fold-change for angiogenic genes** | | | | | | |
|---|---|---|---|---|---|---|---|
| ***ANGPT1*** | 0.048 | 0.420 | ***0.731*** | 0.067 | ***0.631*** | 0.290 | *1.295* |
| ***VEGFA*** | 0.325 | 0.407 | 0.241 | 0.225 | 0.072 | *1.343* | 0.155 |
| ***VEGFB*** | 0.080 | 0.069 | 0.055 | 0.069 | 0.075 | 0.367 | 0.010 |
| | ***A*** | ***B*** | ***C*** | ***D*** | ***F*** | ***G*** | ***H*** |

**Supplemental Table 9: Significance in angiogenic gene expression between donor variability experiments at Day 56.** A value $p<0.05$ indicates significantly different expression between passage schemes p4-5 and p3-5. All $p<0.05$ are red and italicized in the table. Donors A, B, and G are male. Donors C, D, F, and H are female. Donor E was not analyzed in p4-5 due to lack of cells available at cell seeding and therefore not present in this table.

1 out of 3 (33%) angiogenic genes have low variability between passage schemes with fewer than 3 donors (less than 50%) showing significant differences in fold change. This gene name is bolded and italicized in the table: VEGFB.

Donors B (male), C (female), G (male), and H (female) are the most variable between passage scheme: 2 out 3 angiogenic genes have significantly different expression. Donors A (male), D (female), and F (female) are the least variable between passage scheme: 1 out of 3 angiogenic genes has significantly different expression.

| | **p-values for angiogenic gene expression between p4-5 and p3-5** | | | | | | |
|---|---|---|---|---|---|---|---|
| ANGPT1 | 0.709 | *0.012* | *0.014* | 0.286 | *0.021* | 0.087 | *0.027* |
| VEGFA | *0.005* | *0.023* | *0.001* | *3.15E-05* | 0.425 | *1.13E-04* | *0.029* |
| ***VEGFB*** | 0.109 | 0.152 | 0.283 | 0.112 | 0.460 | *0.002* | 0.855 |
| | ***A*** | ***B*** | ***C*** | ***D*** | ***F*** | ***G*** | ***H*** |

**Supplemental Table 10: Order of magnitude differences between passage schemes p405 and p3-5 for donor variability experimental assays at Day 56.** A value of 1 indicates a one order of magnitude (OOM) difference in output between passage schemes p4-5 and p3-5. All values greater than 1 are red and italicized, and all values that round to 1 (i.e., greater than 0.5) are bolded and italicized. Donors A, B, and G are male. Donors C, D, F, and H are female. Donor E was not analyzed in p4-5 due to lack of cells available at cell seeding and therefore not present in this table.

Most experimental outputs differ by less than one OOM between passage schemes at Day 56. Cell number is the most variable output: 1 donor (G, male) has an OOM difference greater than 1 and 1 donor (F, female) has an OOM difference greater than 0.5. ALP activity, calcium fold change, and phosphorus fold change are the least variable outputs: all donors have OOM differences less than 0.5. Donors A (male) and H (female) have OOM differences less than 0.5 for all experimental assays.

| | **Order of magnitude difference for experimental assays between p4-5 and p3-5** | | | | | | |
|---|---|---|---|---|---|---|---|
| ***Metabolic activity*** | 0.206 | ***0.649*** | 0.083 | 0.125 | 0.073 | 0.416 | 0.152 |
| ***Cell number*** | 0.267 | 0.456 | 0.327 | 0.215 | ***0.706*** | *1.315* | 0.141 |
| ***OPG secretion*** | 0.063 | 0.269 | ***0.537*** | ***0.547*** | 0.041 | 0.372 | 0.096 |
| ***ALP activity*** | 0.236 | 0.292 | 0.176 | 0.237 | 0.228 | 0.202 | 0.183 |
| ***Calcium fold change*** | 0.065 | 0.087 | 0.092 | 0.104 | 0.138 | 0.057 | 0.131 |
| ***Phosphorus fold change*** | 0.049 | 0.087 | 0.089 | 0.085 | 0.113 | 0.031 | 0.110 |
| | ***A*** | ***B*** | ***C*** | ***D*** | ***F*** | ***G*** | ***H*** |

**Supplemental Table 11: Significance between passage schemes p4-5 and p3-5 for donor variability experimental assays at Day 56.** A value p<0.05 indicates significantly different outputs between passage schemes p4-5 and p3-5. All p<0.05 are red and italicized in the table. Donors A, B, and G are male. Donors C, D, F, and H are female. Donor E was not analyzed in p4-5 due to lack of cells available at cell seeding and therefore not present in this table.

There are significant differences at Day 56 between passage schemes p4-5 and p3-5 for most assays. There are no significant differences between passage schemes at Day 56 in: metabolic activity for donors D and F (both female), cell number for Donor H (female), OPG secretion for donors A (male), F (female), and H (female), and phosphorus content for Donor G (male).

| | **p-values for experimental assays between p4-5 and p3-5** | | | | | | |
|---|---|---|---|---|---|---|---|
| ***Metabolic Activity*** | *9.27E-05* | *4.44E-06* | *0.009* | 0.194 | 0.053 | *2.50E-06* | *0.004* |
| ***Cell Number*** | *0.015* | *2.59E-04* | *0.001* | *1.92E-04* | *0.047* | *1.87E-04* | 0.585 |
| ***OPG secretion*** | 0.466 | *0.047* | *5.26E-07* | *0.007* | 0.770 | *0.003* | 0.270 |
| ***ALP activity*** | *1.57E-06* | *3.52E-04* | *0.001* | *1.72E-04* | *3.52E-04* | *0.003* | *0.001* |
| ***Calcium Content*** | *0.005* | *1.76E-06* | *3.20E-04* | *0.008* | *3.67E-06* | *0.009* | *1.50E-04* |
| ***Phosphorus Content*** | *0.012* | *2.64E-06* | *4.49E-04* | *0.017* | *5.63E-05* | 0.055 | *0.001* |
| | ***A*** | ***B*** | ***C*** | ***D*** | ***F*** | ***G*** | ***H*** |
